# Detecting phenotype-specific tumor microenvironment by merging bulk and single cell expression data to spatial transcriptomics

**DOI:** 10.1101/2024.02.23.581547

**Authors:** Wencan Zhu, Hui Tang, Tao Zeng

**Author notes:** The corresponding authors should be contacted to Tao Zeng. These authors contributed equally to this work (Wencan Zhu, Hui Tang).

## Abstract

In addressing the limitations of current multimodal analysis methods that largely ignore phenotypic data, leading to a lack of biological interpretability at the phenotypic level, we developed the Single-Cell and Tissue Phenotype prediction (SCTP), a deep-learning-based multimodal fusion framework. SCTP can simultaneously detect phenotype-specific cells and characterize the tumor microenvironment of pathological tissue by integrating essential information from the bulk sample phenotype, the composition of individual cells, and the spatial distribution of cells. Upon evaluating SCTP’s efficiency and robustness against traditional analytical methods, we developed a specialized model, SCTP-CRC, tailored for colorectal cancer (CRC). This model integrates RNA-seq, scRNA-seq, and spatial transcriptomic data to offer a better understanding of CRC. SCTP-CRC has proven effective in accurately identifying tumor-associated cells and clusters and continuously defines boundary regions as well as the spatial organization of the entire tumor microenvironment. This enables a detailed depiction of cellular communication networks, capturing the dynamic shifts that occur during tumor progression. Furthermore, SCTP-CRC extends to the identification of abnormal sub-regions in the early state of CRC and uncovers potential early-warning signature genes such as MMP2, IGKC, and PIGR. These biomarkers are not only important in recognizing the onset of CRC but may also play a crucial role in differentiating between CRC-derived liver metastases and primary liver tumors. SCTP stands as a transformative framework, offering a deeper understanding of the tumor microenvironment through its ability to quantitatively characterize cancer’s fundamental traits and dissect the intricate molecular and cellular interactions at play. This comprehensive insight supports the early diagnosis and enables personalized treatment strategies, marking a significant stride toward improving patient outcomes and tailoring therapies to individual disease profiles.

## Introduction

Tumor complexity is portrayed by the spatial diversity and heterogeneity of cells. Single-cell sequencing technologies are fundamentally transforming biomedical research and clinical applications, and are disclosing detailed intra- and inter-molecular insights into organized cells from healthy and pathological tissues ([1]). The establishment and the rapid growth of single-cell RNA sequencing (scRNA-seq) is becoming an efficient and cost-effective identification of molecular profiles of cells at the individual level rather than at the population level ([2, 3]). However, during the tissue dissociation stage of scRNA-seq, the single cell technology would loss the original spatial positioning of single cells and their closeness within the native tissue ([4]). By contrast, the emergence of spatial transcriptomics (ST) technologies has shown promise in elucidating the spatial structures of tissues, mapping the subcellular localization of mRNA molecules at the cellular level ([5]). ST data typically reflect an average of gene expressions from multiple cells in a given tissue region, complicating the precise characterization of specific cellular spots. ST approaches and data advance the study of developmental biology, and help deepen our knowledge of tumor biology. Especially, ST enables the spatial localization of cells alongside profiling spacial expression patterns of marker genes, therefore provides a unique chance to explore the diverse molecular heterogeneity under the cellular heterogeneity within the tumor development and progression ([6]).

Dissimilar to RNA-seq as bulk data, scRNA-seq and ST data can distinguish cell or spot with different cell types, states, and correlations in a mixed tissue environment ([7, 8, 9]). Cell or spot clusters are typically identified using unsupervised clustering, where each cluster’s marker genes and their correlation with known cell types and pathways are further examined to elucidate each cluster’s biological significance ([10, 11, 12]). However, unsupervised clustering of cells or spots would fall short in recognizing specific subpopulations with key phenotypes, such as tumor development, invasion and treatment response. This under-estimated information is critical because it should help detect prognostic markers and treatment targets that are particular to individual tumor cells or spots ([13, 14]). On the other hand, although scRNA-seq and ST resolution enable thorough cataloging of cancer subpopulations and adjacent niche cell subpopulations, their clinical application is hindered by the significant costs and labor required for sequencing large cohorts ([15]). The current research landscape shows a significant gap in phenotype prediction at the cellular or spot level, primarily due to insufficient integration of key data information: phenotype information from RNA-seq, cellular composition from scRNA-seq, and spatial details from ST data. Despite the potential for improved prediction accuracy through data synthesis, this area remains underexplored in the scientific community.

Indeed, bulk sequencing data can provide clinical phenotype details of a large number of cohorts, which are widely accessible from the public resources like Gene Expression Omnibus ([16]) and The Cancer Genome Atlas (TCGA ([17])). These data sets provide tumor information about cancer sites, occurrence, progress and treatment response ([18, 19]). Bridging phenotypic bulk data, it should open a new avenues to predict cell / spot phenotypes for high resolution single cells and spatial spots. Recently, scRNA-seq based methods such as Scissor ([20]) and scAB ([21]) have advanced our understanding of cell subpopulations in relation to specific phenotypes measured on bulk samples. These methods are built dependent on the expression correlation between scRNA-seq and RNA-seq data. Scissor employs a regularized regression model for selecting phenotype-relevant cells, and scAB uses a matrix factorization model to infer cell states linked with individual phenotypes. These methods focused on the association inference between single cells and bulk samples, rather than prediction of cellular phenotypes. Meanwhile, scATOMIC ([22]) represents a comprehensive pancancer tool for classifying malignant and non-malignant cell employing random forest models. However, it is only based on single-cell data without phenotypic information from bulk samples. On the other hand, in the realm of spatial transcriptomics, SpaRx ([23]) and Cottrazm ([24]) are emerging recently for phenotype prediction task. SpaRx predicts cellular reactions to drugs by transfer learning from drug responses of cell lines, and Cottrazm identifies the tumor boundary connecting malignant and non-malignant cell spots in tumor tissues through integrating ST with hematoxylin and eosin histological image. The two methods predict phenotypes of spatial spot by integrating spatial gene expression profiles with bulk or image data, take into consideration the cell compositions in spatial spots. As of now, there are still great gaps in integrating ST with scRNA-seq and RNA-seq data. This capability is particularly crucial in the context of examining cellular heterogeneity within the tumor microenvironment since accurate identification of malignant regions not only offers diverse biological insights of tumor dynamics but also is instrumental for personalized medicine in clinical application.

Motivated by these challenges and requirements, we developed a new machine learning approach to predict cell and spot phenotypes, called Single-Cell and Tissue Phenotype prediction (SCTP). It combines paired or unpaired bulk RNA sequencing (RAN-seq), single-cell RNA sequencing (scRNA-seq) and spatial transcriptomics (ST) data, where the phenotype information can be obtained from bulk samples (Figure 1A). To model the tripartite relationship among the three aforementioned data sources, the SCTP framework is structured to initially assess the correlations between individual cells and bulk samples, as well as between spots and bulk samples Concurrently, it integrates cell-cell and spot-spot networks to represent the correlations between individual cells and spots, as depicted in Figure 1B. Following this initial assessment, SCTP employs a deep multi-task fusion learning approach to construct a predictive model for phenotypes. This model leverages the quantitatively identified phenotype associations for cells and spots to characterize the temporal-spatial distribution of molecular and cellular phenotypes across scales—from individual cells, through spatial regions, to entire samples, as shown in Figure 1C. After comprehensively evaluating the efficiency and robustness of the model, SCTP has been applied to develop a microenvironment landscape for colorectal cancer (CRC), termed SCTP-CRC, through the integration of various omics datasets pertaining to CRC and its liver metastasis. SCTP-CRC demonstrates its efficacy in accurately identifying tumor-associated cells, their clustering and the delineation of boundary regions. This capability further encompasses the detection of early-stage abnormal sub-regions in colorectal cancer (CRC), thereby enabling the identification of novel early-warning signature markers. The spatial components and organizations of the tumor microenvironment largely reflect the cellular communication network, which is consistent with the dynamics observed in tumor progression. This consistency is supported by the spatial patterns of marker gene expression, functional activities, and cellular composition. Notably, many of these marker genes (such as MMP2, IGKC, PIGR) may play analogous roles in liver metastases from CRC, as opposed to primary liver cancer, supporting the specificity and sensitivity of SCTP-CRC findings.

**Figure 1:**
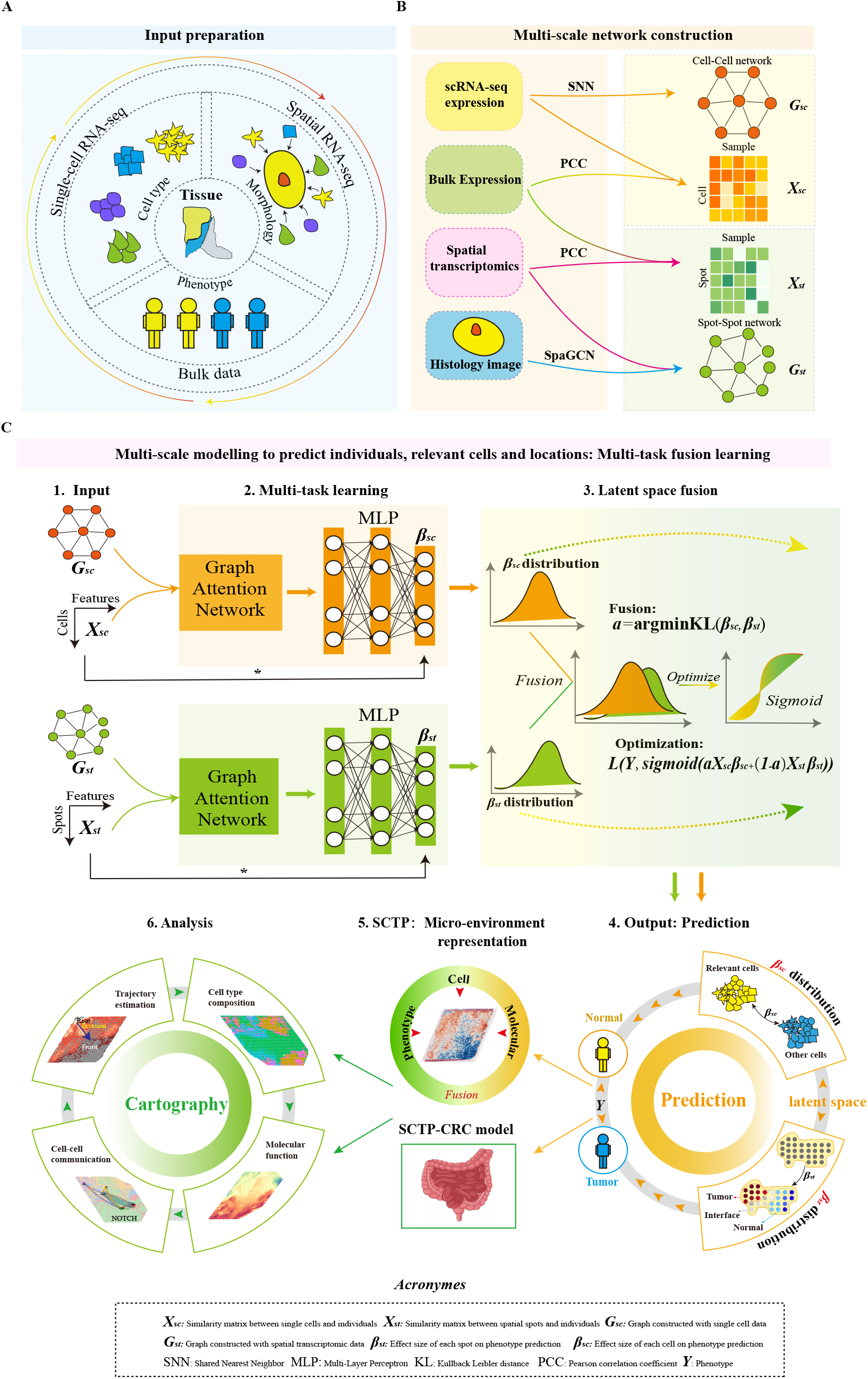
Workflow of SCTP. **A** SCTP necessitates five inputs: a single-cell expression matrix, a spatial transcriptomics matrix, spots location matrix, pathology image and a bulk expression matrix accompanied by corresponding sample phenotypes. **B** SCTP calculates a correlation matrix and a cell–cell similarity network based on single-cell and bulk data, similarly a correlation matrix and a spot–spot similarity network based on spatial transcriptomics and bulk data. **C** Detailed SCTP analysis framework. consists of six parts: The constructed graphs and feature martrix as data inputs. SCTP estimates cell and spot effects through a multitask fusion learning framework including multi-task learning layer and latent space fusion layer. The coefficients were then integrated for phenotype prediction. The macro-phenotype from bulk data are transferred to micro-phenotype of cells and spots and SCTP-CRC is a showcase to demonstrate the functionality of SCTP. Downstream analysis can be carried out following SCTP’s prediction.

Our SCTP methodology provides a valuable approach to analyze and understand the tumor microenvironment from an innovative and integrative perspective by combining the essential information from the bulk sample phenotype, single cell composition and cellular spacial distribution. As an automated model for tissue phenotype prediction, SCTP facilitates a more profound understanding of tumor microenvironments, enables quantitative characterization of cancer hallmarks, and elucidates the underlying complex molecular and cellular interplay. This machine learning model and analysis tool has great potential for early detection and personalised treatment of colorectal cancer and other complex diseases.

## Result

### SCTP outperforms existing single modality based methods

To evaluate the benchmark performance of SCTP compared to existing methods, we first implemented a single-modality model of SCTP, specifically employing scRNA-seq data integrated with bulk RAN-seq data (Methods). In two different application scenarios, SCTP was extensively compared with two state-of-the-art methods, scAB and Scissor. Of note, a detailed summary of all datasets utilized in our study is provided in Supplementary Table 1 (bulk data 1-8), 2 (SC data 1-4) and 3 (ST data 1-5) including the accession number and sample information of each data.

In one scenario, its objective was to identify cell subpopulations associated to tumor state, where the phenotype is defined as primary CRC vs. Normal. The SCTP is learned respectively on two distinct scRNA-seq data, SC data 2 ([25]) and SC data 3 ([26]), integrated with the bulk data 1 ([27]). In the other scenario related to immunotherapy response, it aims to identify cell subpopulation responsive to immune checkpoint blockade (ICB), and SCTP is built on SC data 1 utilizing phenotypic indicators from melanoma samples (bulk data 6) subjected to treatment regimens including anti-PD-1 monotherapy or a combination therapy of anti-PD-1 and anti-CTLA-4 [28], where individual cells are previously categorized into nine distinct cell types [29].

First, in qualitative assessment, SCTP tends to detect phenotype-specific cells without cell type preference. As the phenotype-specific cells accentuated in red shown in Figure 2A, scAB tends to select more myeloid cells in CRC1, epithelial cells in CRC2 and Endo and Mal cells in CRC3. Scissor also tends to select more myeloid and epithelial cells in CRC2. In contrast, SCTP appears to select high-ranking cells from different types, and would thus be able to find phenotype-specific cells with a more balanced ratio between cell types or states, which is likely to be important for characterizing the tumor region in downstream analysis. The composition of selected cells can be found in Supplementary Figure 1A.

**Figure 2:**
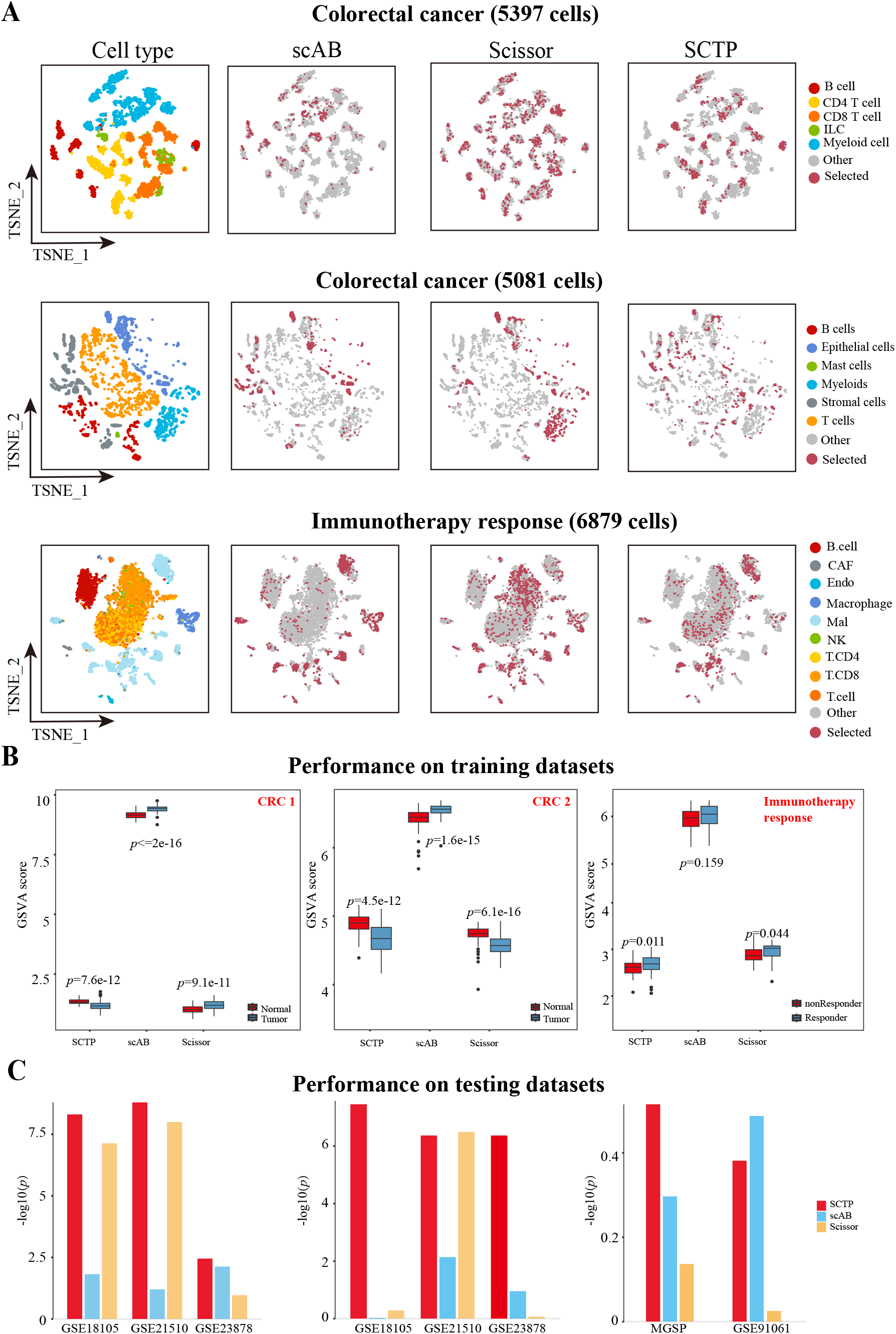
Benchmark test for SCTP in absence of spatial data. **A** (t-distributed Stochastic Neighbor Embedding) (TSNE) visualization of cells from two colorectal cancer or one melanoma microenvironment. The left plot shows the original cell type distribution. Cells identified through different methods (scAB, Scissior and SCTP) are displayed in separated graphs, where red signifies selected cells and grey denotes the remaining cells. **B** Comparisons of GSVA scores calculated using selected cells’ gene signatures between tumor v.s. normal (colorectal cancer) or response v.s. non-response (immunotherapy response). Data are presented as boxplots (minima, 25th percentile, median, 75th percentile, and maxima). **C** The performance of biomarkers on testing datasets, as inferred from different methods, is compared in terms of phenotype differentiation, using P-values from Wilcoxon tests.

Next, in quantitative evaluation, the differentially expressed genes (DEGs) were characterized as genes exhibiting significantly different expressions between phenotype-specific cells and others in accordance to previous assessment work from [21]. Based on these DEGs, the Gene Set Variation Analysis (GSVA) scores for each sample within bulk datasets were calculated to distinguishing two sample groups with distinct phenotypes. According to the comparison significance (e.g. P values determined via Student’s t-test) depicted in Figure 2B, SCTP exhibited higher significance (lower P-value) compared with other methods in the two bulk datasets, bulk data 1 and 3, suggesting that SCTP can efficiently detect phenotype-specific cells, and provide clinically relevant DEGs for explaining bulk data with a feedback way.

Further, to assess the generalization capability of identified cells with predicted phenotype, the GSVA scores were calculated and evaluated in independent validation datasets, which include CRC related data (bulk data 5 ([30]), bulk data 6 ([31]) and bulk data 7 ([32]) ; and immunotherapy-response related data (Melanoma Genome Sequencing Project (bulk data 2) and bulk data 3 ([33])). Compared to Scissor and scAB, SCTP consistently exhibited the lowest P-values on all CRC related datasets, and comparable performances on immunotherapy-response related data (Figure 2C). These results demonstrated that SCTP is capable of precisely identifying cells most strongly associated with a given phenotype when applied to single-cell data in a single-modality model. Of note, the accuracy of SCTP predictions has been comprehensively validated through a 10-fold cross-validation along with a comparison of computational costs among various methods (Supplementary Figure 1B and 1C), demonstrating SCTP’s superior efficiency particularly in terms of reduced time requirements for computation.

### SCTP improves the analysis robustness and stability of phenotype prediction by merging bulk and single-cell expression data to spatial transcriptomics

SCTP not only has its ability to effectively identify phenotypic cells using only single-cell data as scAB and Scissor by the coefficient *β*_*sc*_ in latent fusion space, but also provide robust and stable multi-modal fusion to incorporate ST data. For comparing and evaluating SCTP performance with multi modality fusion, four distinct ST datasets (P1-P4) from ST data 1 in Table 3 ([34]), with encompassing histological information, were integrated into the above CRC case. Both scRNA-seq data (SC data 3 in Table 2) and these four ST datasets were derived from patients with primary colorectal cancer.

For each ST dataset, the SCTP was similarly applied during the training phase, integrating bulk data 1 and SC data 3. The two coefficients *β*_*sc*_ and *β*_*st*_ were estimated in latent fusion space (Methods). Based on the coefficient *β*_*sc*_, cells associated with the specific phenotype are selected (Methods). Subsequently, DEGs are identified by comparing the selected phenotype-associated cells with the remaining cells. Then based on these DEGs, the GSVA scores for each sample in the bulk data 1 were obtained as shown in Figure 3A, allowing for a comparison of the model’s performance with the inclusion of different ST datasets or without. From these results, SCTP demonstrates that the multi-modality method (e.g. SCTP P1 or SCTP P4) are more effective than single-modality method (e.g. SCTP SC), and it is also stable for application of different ST datasets. Subsequently, GSVA score was calculated on bulk samples with the same DEGs from three external validation dataset (bulk data 2, 3 and 4 from Table 1), each comprising both CRC and normal samples. As illustrated in Figure 3B, the GSVA scores effectively discriminate between different phenotypes across all independent bulk datasets. These overall significant results underscore the stable and robust performance of SCTP on integrating bulk and single-cell data to ST data.

**Table 1:** Bulk data information.

|  | Index | Data accession | Sample size | Function | Section |
| --- | --- | --- | --- | --- | --- |
| Colorectal cancer | 1 | GSE44076 | 98 tumor + 98 normal | Training set | SCTP-CRC model |
|  | 2 | GSE18105 | 17 tumor + 17 normal | Validation set | Latent space Evaluation |
|  | 3 | GSE21510 | 19 tumor + 25 normal | Validation set | Latent space Evaluation |
|  | 4 | GSE23878 | 35 tumor + 24 normal | Validation set | Latent space Evaluation |
|  | 5 | TCGA-COAD | I (81), II (167), III (123), IV (66) | Validation set | Biomarker Validation |
| Immunotherapy response | 6 | PRJEB23709 | 46 responder + 27 non responder | Training set | Latent space Evaluation |
|  | 7 | MGSP | 26 responder + 25 non responder | Training set | Latent space Evaluation |
|  | 8 | GSE91061 | 13 responder + 15 non responder | Training set | Latent space Evaluation |

**Table 2:** Single-cell RNA-seq data information.

|  | Index | Data accession | # cells in total | # meta-cells for training | Celltype composition | Section |
| --- | --- | --- | --- | --- | --- | --- |
| Immunotherapy response | 1 | GSE115978 | 6879 | 6879 | B cells, fibroblasts, endothelial cells, macrophages, malignant cells, Natural Killer cells, CD4+ T cells, CD8+ T cells, and T cells | Latent space Evaluation |
| Colorectal cancer | 2 | GSE146771 | 43817 | 5397 | B cells, CD4 T cells, CD8 T cells, ILCs and myeloid cells | Latent space Evaluation |
|  | 3 | GSE132465 | 63689 | 5081 | Epithelial cells, T cells, B cells, myeloid cells, stromal cells, and mast cells | Latent space Evaluation and prediction |
|  | 4 | GSE225857 | 41892 nonimmune cells<br>196473 immune cells | 4100 immune cells | nonimmune cells: tumor cells, fibroblasts, and endothelial cells; immune cells: T cells, natural killer (NK) cells, B cells/plasma cells, monocytes/macrophages, dendritic cells, and mast cells | SCTP-CRC model and prediction |

**Table 3:** Spatial transcriptomic data information.

|  | Index | Data accession | Label | # spots | Pathology segmentation | Section |
| --- | --- | --- | --- | --- | --- | --- |
| Colorectal cancer | 1 | DRA015288 | P1-P4 | P1(2273),<br>P2(2465),<br>P3(2002),<br>P4(2475) | Only on P1 | Latent space Evaluation |
|  | 2 | GSE225857 | Q1-Q2 | Q1(2055),<br>Q2(3418) | ✓ | SCTP-CRC model |
|  |  |  | Q3-Q4 | Q3(3306),<br>Q4(4017) | ✓ | Prediction validation |
|  | 3 | https://zenodo.org/records/755346 | S1-S2 | S1(4992),<br>S2(4992) | ✓ | Prediction external validation |
| CRC-liver metastases | 4 | GSE225857 | L1-L2 | L1(4673),<br>L2(4797) | ✓ | Transformative Prediction |
| Hepatocellular carcinoma | 5 | http://lifeome.net/supp/livercancer-st/data.htm | - | 4992 | ✓ | Biomarker Validation |

**Figure 3:**
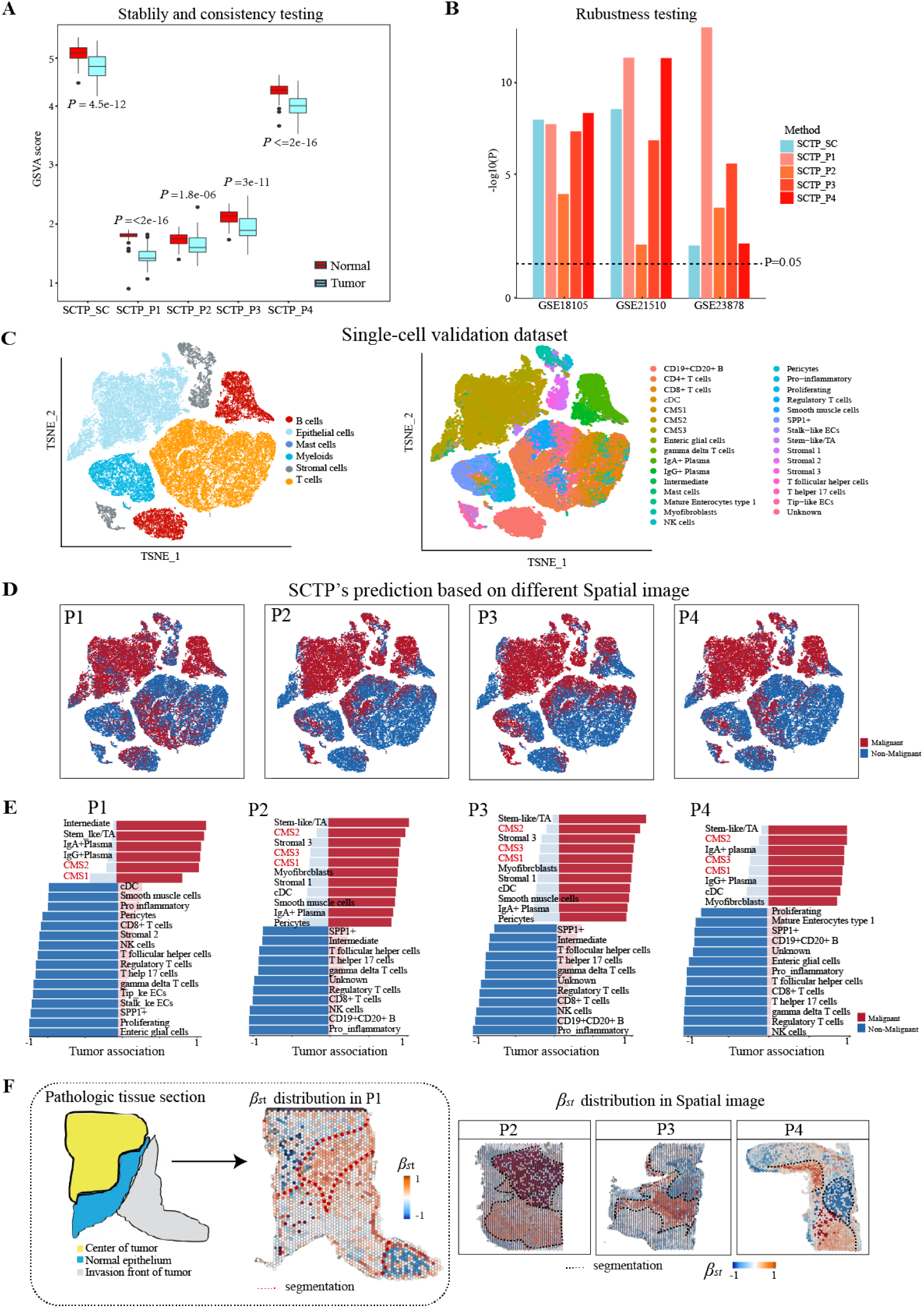
SCTP performance when integrated with scRNA-seq and spatial transcriptomic data. **A** Comparisons of GSVA scores calculated using selected cells from models integrating different spatial transcriptomic datasets(minima, 25th percentile, median, 75th percentile, and maxima). **B** Comparison of SCTP with different ST data based on the predictive performance on three external testing datasets by using P-values from Wilcoxon tests. The dashed black line indicates a p-value=0.05 **C** TSNE representation of single cell database used for prediction. The cells are colored by global cluster and subcluster annotations respectively. **D, E** Cell phenotype (malignant v.s. non-malignant) prediction based on different spatial transcriptomic datasets by SCTP. **D** TSNE presentation, cells are colored by predicted phenotype. **E** Partition of predicted malignant and non-malignant cells in each cell subtypes. The estimated *β*_*sc*_ is between -1 and 1. **F** Estimated *β*_*st*_ in latent space mapping to the histology images. Regions with opposite signs on *β*_*st*_ are segmented with dotted line.

### SCTP provides consistent prediction of phenotype-associated cell subpopulations across ST datasets

The principal utility of SCTP lies in its capacity to precisely predict the phenotype of individual cells and specific spatial spots. Thus, we further analyzed SCTP prediction on SC data 3 that included six primary global types: T cells, B cells, Epithelial cells, Mast cells, Myeloid cells, and Stromal cells; and each of these categories can be subdivided into various subtypes, reflecting the heterogeneity inherent in cellular structures and functions (Figure 3C). For example, the epithelial cell includes colorectal cancer (CRC) subtypes—namely CMS1, CMS2, and CMS3—as defined by the Consensus Molecular Subtypes (CMS) classification ([35]). Each CMS subtype exhibits distinctive characteristics: CMS1 is marked by pronounced immune activation; CMS2, referred to as the “canonical” subtype, generally presents a favorable prognosis and is the most prevalent; CMS3 is distinguished by metabolic dysregulation.

To evaluate the coefficient estimation of latent space and prediction accuracy of SCTP, we utilized the 5081 meta-cells of SC data 3 from Table 2, and train the SCTP model employing different spatial datasets in ST data 1 from Table 3. Following the training phase, we then utilized these fine-tuned models to predict the phenotype of the remaining cells in SC data 3 as validation using Equation (9).

Using different ST datasets (P1-P4), SCTP was able to predict cell states as malignant or non-malignant depicted in Figure 3D. These visual representations reveal a predominance of epithelial cells classified as malignant; in contrast, other cell types are predominantly predicted as non-malignant. Further exploration of these predictive results was undertaken, and cell subtypes with the highest proportion (over 70%) of cells predicted as malignant or non-malignant are concentrated (Figure 3E and Supplementary Data 2), such as a predominance of CMS1, CMS2, and CMS3 cells among those predicted to be tumor cells. These alignments not only validated again the predictive accuracy and consistency of SCTP, but also underscored the critical role of CMS classification in understanding and predicting tumor behavior. An additional intriguing aspect that emerged from the analysis was the predominance of immune cells, specifically T cells and B cells, along with the malignant cells. This finding opens up paths for further investigation into the function of immune cells in tumor suppression and the potential for leveraging this understanding in developing targeted therapies ([36]).

### SCTP enhances the division of tumor microenvironment by quantitative phenotype indicator

According to those stable and robust evidences, SCTP indeed have better performance on single-cell representation compared to conventional methods, and its another methodology merit would be simultaneously obtain the cell spatial representation. For the same SC data 3 and ST data 1 as above mentioned, we further analyzed SCTP prediction of coefficient *β*_*st*_ to distinct spatial spots (Methods section) for the segmentation of the pathological tissue (Figure 3F). Compared to the pathological tissue segmentation previously reported [34] as displayed on the left panel of Figure 3F, the estimated coefficient *β*_*st*_ superimposed on the histological data revealed that the spatial effects estimated by SCTP align closely with the boundaries delineated in the pathological segmentation. This case (SCTP P1) particularly highlighted that the upper-left subregions of the histological data as identified by SCTP show a positive association with the tumor phenotype, suggesting a high probability that these areas should be predominantly composed of tumor cells or tumor associated cells (e.g. CAF). Conversely, the subregions characterized by negative *β*_*st*_ values, indicate a reduced likelihood of being primarily tumorous.

For the other three remaining histological datasets (P2-P4), SCTP locates subregions that show clear tumor characteristics (Figure 3F), although their corresponding pathological tissue segmentation is not available for evaluation. Critically here, SCTP has demonstrated the ability to discern tumor-associated areas even in the absence of pathological benchmarks, especially in contexts where reference material is unavailable. Additionally, we found that integrating single-cell data can actually help produce more precise segmentation results. When ST data without scRNA-seq data is used as the single modality model of SCTP, the estimated coefficients yield a visually less distinct segmentation of tumor regions, the observation of which is consistent across all four datasets (P1-P4) as in Supplementary Figure 2 shown.

### SCTP-CRC demonstrates precise phenotypic partitioning of CRC tumor microenvironment

To demonstrate the systematic analytical capabilities of SCTP, we conducted an indepth study of the tumor microenvironment in colorectal cancer (CRC), utilizing the large scRNA-seq and ST data recently available for CRC. We have built a CRC-specific SCTP model (i.e. SCTP-CRC), based on the same bulk data 1 from Table 1, SC data 4 from Table 2 and ST data 2 from Table 3 which collected four samples of primary CRC patients (labeled as Q1-Q4). The addition of pathological diagnoses from this ST data serves as a crucial reference, allowing for a more precise assessment on spatial segmentation. We first trained SCTP-CRC using a selected subset of immune meta-cells together with the ST dataset Q1 or Q2. Coefficients *β*_*sc*_ of each cell and *β*_*st*_ of each spot were estimated as described in Methods section. As shown Figure 4A, the estimated values 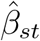 from Q1 and Q2 indicate the relationship between spots and phenotype with clear tumor region segmentation globally aligned to pathological segmentation. Results on Q3 and Q4 can be found in Supplementary Figure 3A.

**Figure 4:**
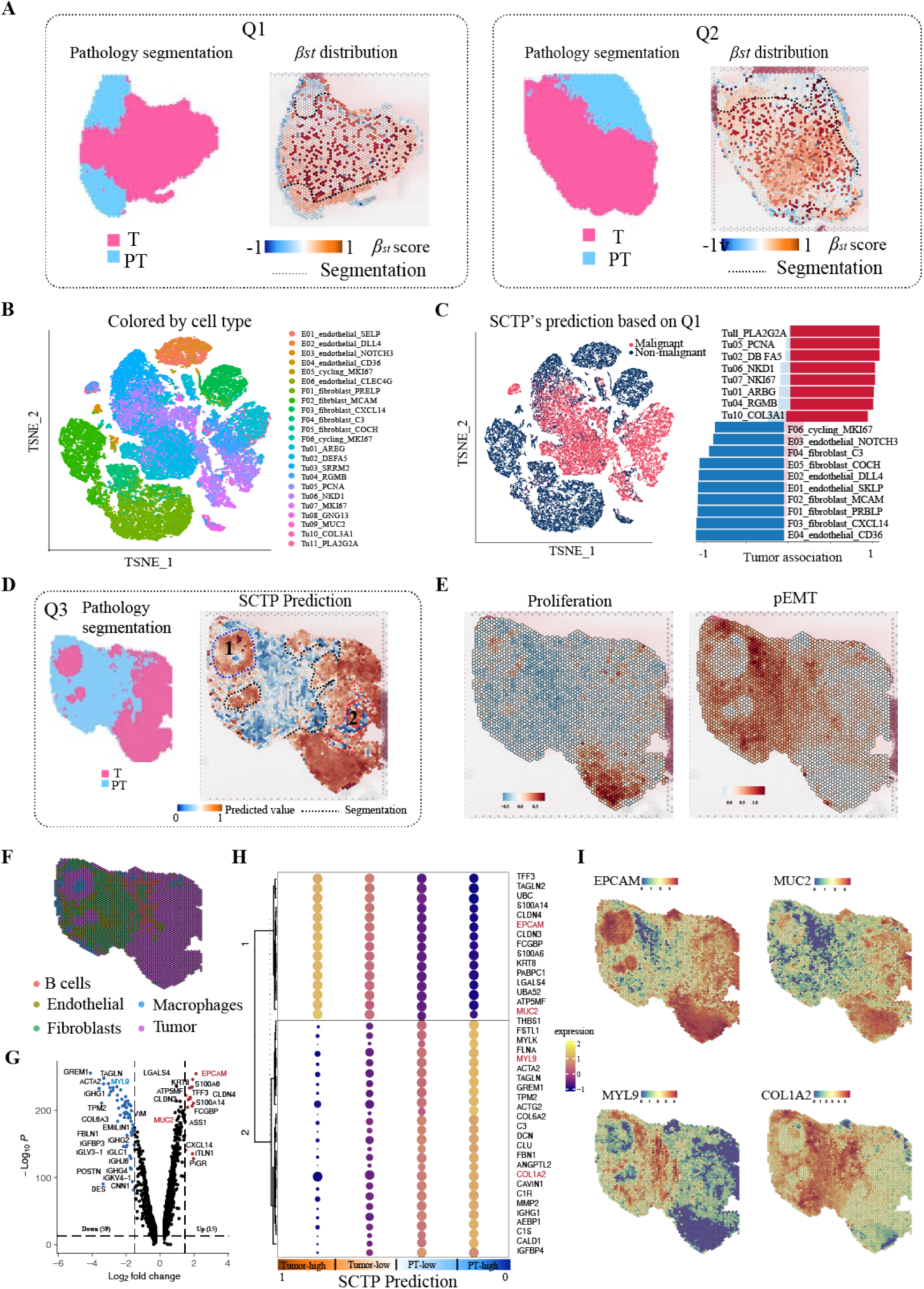
SCTP model of colorectal cancer (SCTP-CRC model) and its prediction of tumor microenvironment partitioning. **A** Pathology segmentation and estimated *β*_*st*_ in latent space for Q1 and Q2 respectively. Regions with opposite signs on *β*_*st*_ are segmented with dotted line. **B** TSNE representation of 41892 non-immune cells, colored by original cell type annotations. **C** Prediction of malignancy of the non-immune cells based on the SCTP-CRC model. Left: TSNE presentation as in **B**, cells are colored by predicted phenotype. Right: the partition of predicted malignant and non-malignant cells in each cell type. **D** Pathology segmentation and prediction on Q3 histological image based on the SCTP-CRC model. **E** Expression level of gene lists in the pathways related to Proliferation and Epithelial-Mesenchymal Transition (EMT) respectively. **F** Estimated cell types in each spot, which is identified as the most dominant cell cluster. **G** Volcano plot showing differentially expressed genes between most tumor alike spots and least tumor alike spots. **H** DEGs between different groups defined by predicted tumor likelihood. **I** Feature plots showing expression level of EPCAM, MUC2, MYL9 and COL1A2 as identified from **H**.

For clarity, the term ’SCTP-CRC’ model is herein used to refer specifically to the model developed using ST data from dataset Q1 as an example for illustration. This model was subsequently applied to various downstream analytical tasks. As shown in Figure 4B, SCTP-CRC can predict the phenotype (malignant cell or non-malignant cell) of tumor cells, endothelial cells, fibroblasts and others. In fact, SCTP-CRC assigned each cell a predicted probability of being a tumor one, where the likelihood is calculated using Equation 9. According to the prediction, SCTP-CRC has demonstrated remarkable accuracy in identifying tumor cells (Figure 4C), especially notable in tumor cells expressing specific marker genes such as PLA2G2A, PCNA, and NKD1. Impressively, over 90% of cells marked by these genes were precisely classified as malignant, and the relevance of most marker genes in the context of CRC has been previously established ([37, 38, 39]). This finding underscores the effectiveness of SCTP in discerning tumor cells based on immune cell profiles, and suggests that the immune cell profiles within the same tumor microenvironment can be predictive of the properties of non-immune cells. This indicates a broader suitability of SCTP in deciphering complex cellular interactions in cancerous tissues, thereby providing substantial insights for advancements in cancer diagnostics and research.

Next, we proceeded to analyse the SCTP-CRC prediction on ST dataset Q3. According to a comparative observation against pathologically defined segmentation shown in Figure 4D, there is actually a high degree of concordance between the predicted and actual tumor subregions, indicating the accuracy of SCTP-CRC for tumor region identification. Considering above molecular heterogeneity in the spatial dimension, we next explored the functional heterogeneity in tumor microenvironment by several signature scores (in Figure 4E and Supplementary Figure 3C), particularly the association with cellular proliferation ([40]) and Epithelial-Mesenchymal Transition (EMT). Notably, elevated proliferation scores, indicative of active tumor growth and division, were predominantly observed in the lower-right regions of the tissue, suggesting these areas as tumor-microenvironment hotpots. These scores also highlighted peripheral regions, particularly the margins of a smaller tumor area located in the upper-left section, suggesting dynamic biological activities at these tumor margins ([41]). Meanwhile, EMT scores are also important indication in tumor invasion, as EMT enables cancer cells to acquire mobility and invasiveness ([42]). High expression in the tumor surrounded region (as shown in right panel of Figure 4E) indicated that two small subregions on the left of the image have an increased activity of the EMT procedure in the cells located at the periphery of the tumor. Such upregulated EMT facilitates the detachment of cancer cells from the primary tumor mass, enabling them to invade surrounding tissues [43]. As known, tumors are often heterogeneous, containing regions with varying degrees of EMT, which can be actually observed that different EMT scores quantified different tumor associated regions. The tumor margin regions, showing differed EMT activity, can exhibit distinct molecular and cellular characteristics compared to the tumor core regions. Additional signature scores also supported these observations and conclusions, including cell cycles pathways and tumor evasion related gene sets (Supplementary Figure 3C). According to the spatial map of cellular composition from deconvolution ([44]) using single cell reference from the same study of [45] (Methods and Supplementary Table 1), the dominant cell type in each spot where subregions predominantly occupied by tumor cells are depicted in purple (Figure 4F). The SCTP-CRC demonstrated high accuracy in predicting cellular distributions within tumor regions, and achieved a high degree of global alignment with results obtained from pathological segmentation and deconvolution analyses. Of note, SCTP-CRC was more effective in deriving high-resolution distributions of non-malignant cells across the examined tumor landscapes compared to deconvolution results. For instance, although both SCTP-CRC and deconvolution approaches identified tumor region 1 in Figure 4D, the annotations from SCTP-CRC exhibited greater precision in detecting a small, distinct non-tumor region within tumor region 2, clearly separating it from its surroundings.

Furthermore, we compared the gene expression discrepancies between predicted tumors and peritumoral (PT) subregions, gaining biological significance of predicted tumor spots. Each spot comprises cells potentially originating from various types, so that, the SCTP prediction for each spot is formulated as a continuous probability. We categorized the spatial spots into four groups, differentiated by their predicted probabilities of being tumor-alike. Subsequently, we identified DEGs between the group with the highest probability of being a tumor (prediction *>* 0.5) and the group with the lowest probability (prediction *<* -0.5). This analysis led to the identification of 25 up-regulated and 52 down-regulated genes, as shown in Figure 4G. A significant correlation was observed between the predicted tumor likelihood and the expression levels of these DEGs (Supplementary Data 3), as illustrated in the middle part of Figure 4H (and Supplementary Data 3). This finding reinforces the predictive accuracy of SCTP-CRC and provides valuable realizations into the gene expression profiles associated with phenotypes of tumor spots. Notably, among the up-regulated genes in top-ranking DEGs (Figure 4I), many genes, such as S100A6 and EPCAM, were reported to be associated with the regulation of CRC ([46, 47]). Meanwhile, many down-regulated DEGs, including several IGH (immunoglobulin heavy chain) genes, are indicative of the body’s immune status or response to the cancer. The expression pattern of IGH genes can contribute to specific immune signatures in CRC, which may be associated with CRC prognosis. For instance, a robust B cell response might suggest an active immune environment and potentially better outcomes ([48]).

### SCTP-CRC detects early molecular and cellular signal of colorectal cancer

We made some interesting discoveries from SCTP-CRC predcition on another ST dataset Q4 (Figure 5A). Referring to the histological image, we can segment this image into three distinct subregions, reffered to as subregion 1 to 3. Apparently, the SCTP-CRC predictions for tumor regions agree well with the pathological classifications in subregion 3. Interestingly, a notable observation occurred in subregion 2, which was identified as tumorous in SCTP-CRC but not in the original pathologic evaluation. This discrepancy highlights a region of interest (ROIs) for further investigation and underscores the complexity of tumor conditions.

**Figure 5:**
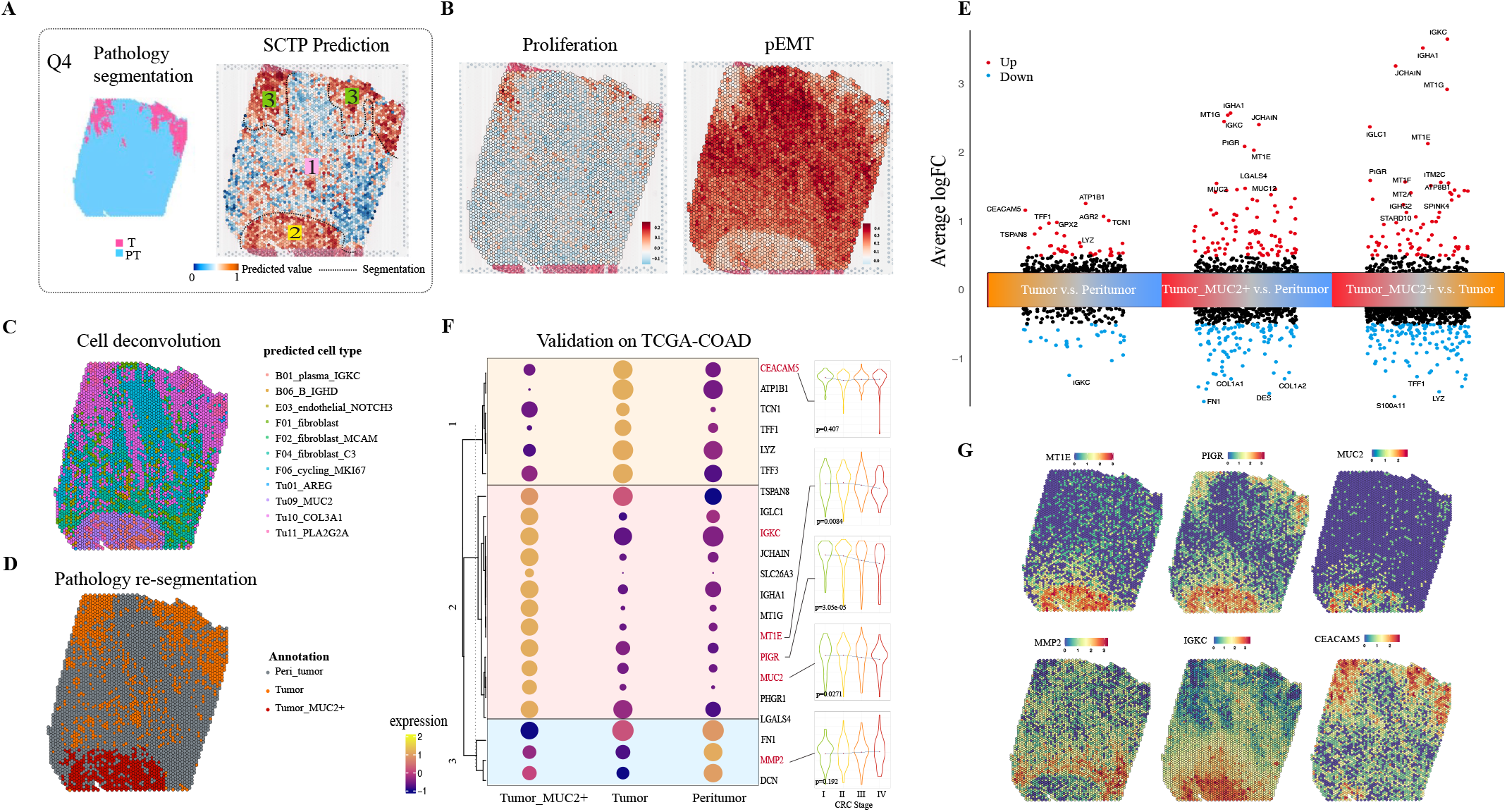
SCTP-CRC detects early signal of colorectal cancer on cell states and spots distribution levels that were unseen in traditional pathological diagnoses. **A** Pathology segmentation and prediction on Q4 histological image based on the SCTP-CRC model. **B** Expression level of gene lists in the pathways related to Proliferation and Epithelial-Mesenchymal Transition (EMT) respectively. **C** Cell composition which shows the most dominant cell cluster. **D** Division of predicted tumor spots into two areas: Tumor as identified in pathological segmentation and Tumor MUC2+ as not shown in pathological segmentation. **E** Volcano plot showing differentially expressed genes between the groups defined in **D. F** The dot plot showing expression level of DEGs identified in **E**, and their expression in TCGA-COAD cohort with different CRC stages. **G** Feature plots showing expression level of genes MT1E, PIGR, MUC2, MMP2, IGKC and CEACAM5.

Actually, subregion 2 was similar to subregion 3, and had enrichment of signature score with EMT pathway (Figure 5B); meanwhile, subregion 3 also revealed predominant signature score with proliferation pathway. Given subregion 3 represents an tumor exhibiting a more advanced developmental pattern with pathological diagnose evidence, we would hold an assumption that subregion 2 might represent an early tumor state of CRC with hidden developmental pattern in clinic viewpoint. Combined with the deconvolution results ([45])(Methods and Supplementary Table 1), subregion 2 would be mainly composed of B cells, fibroblasts and tumor cells (Figure 5C). Considering the subregion 2 exhibited specificity cell type annotated as Tu09 MUC2, therefore it was defined as Tumor MUC2+ for downstream analysis and discussion. Consistent to pathological diagnosis, subregion 3 was defined as Tumor and subregion 1 was defined as Peri-tumor (Figure 5D). To further investigate the molecular difference between the three different areas, DEGs were identified between each pair of them (Figure 5E and Supplementary Data 5). When comparing Tumor to its surroundings peri-tumor, we can find the well-studied cancer genes (Figure 5E). CEACAM5 is over-expressed in nearly 70 percent of epithelial malignancies including colorectal cancer (CRC) ([49]) and has lead to the growth of a novel antibody–drug for the treatment of CEACAM5-positive Epithelial Tumors ([50]). TFF3 is recently demonstrated to be a promoter of clonogenic survival in CRC cells ([51]). Many genes are particularly upregulated DEGs detected in Tumor MUC2+, including genes associated with the immune system such as IGKC and IGHCA1, and genes related to heavy metal processing such as MT1G and MT1E.

To validate our hypothesis concerning the role of Tumor MUC2+ in the early stages of tumor development, we undertook a detailed analysis of differentially expressed genes (DEGs) in the Tumor MUC2+ area, as illustrated in Figure 5F and detailed in Supplementary Data 6. These DEGs represent the most significantly altered genes between the Tumor and Tumor MUC2+ regions. Their expression levels were then cross-referenced with data from the TCGA-COAD cohort (The Cancer Genome Atlas-Colon Adenocarcinoma), which included 435 colorectal cancer patients with four different grades (I to IV). Our analysis revealed that certain genes, including MUC2, MT1E, and PIGR, exhibited higher expression levels in patients at stages I/II, with a noticeable decrease in expression as the disease advanced to more advanced stages, as depicted in Figure 5G. The significance of these trends was determined through p-values obtained from linear regression analyses, which assessed gene expression patterns across successive cancer stages. In contrast, markers prominently expressed in the Tumor area, such as CEACAM5, did not demonstrate significant expression variance across the stages within the TCGA-COAD cohort. These findings lend substantial support to our key hypothesis, suggesting that these key DEGs may serve as early-warning indicators in the early stage of CRC development.

Specifically, within the spatial context, the genes MMP2 and DCN are only highly expressed in area surrounding Tumor MUC2+ but not around Tumor (Figure 5G and Supplementary Figure 3D). MMP2 facilitates tumor invasion and metastasis through ECM degradation ([52]), while DCN acts as a tumor suppressor, modulating growth factor activities and ECM structure ([53]). The upregulation of MMP2 may suggest the process of ECM degradation to create a pathway for further tumor expansion and invasion. Simultaneously, the increased expression of DCN could reflect a response to the changing microenvironment, necessitating the remodeling of the ECM to accommodate tumor growth invasion. Our observed overexpression of both genes at the tumor margin indicates a dynamic interaction through various molecules and signaling pathways.

Besides, multiple immune system related genes are upregulated in Tumor MUC2+, including IGKC, IGHA1 and PIGR, as shown in Figure 5F and Supplementary Figure 3D. The strong association of these genes with the activity of B cells and the humoral immune response indicated that Tumor MUC2+ should be at an early state of CRC ([54]) [55]. Moreover, multiple Metallothionein (MT) genes (MT1A, MT1E, MT1F and MT1G) are also upregulated in Tumor MUC2+ (Figure 5E and Supplementary Figure 3D), and metallothioneins are known to be a family of proteins that play a significant role in metalion homeostasis and detoxification. MT1G has been discussed in [56] for its role in immune filtration in colorectal cancer, where patient with positive MT expression have a poor survival [57]. Understanding the regulation of these genes in different tumor areas (e.g. Tumor MUC2+ and conventional Tumor here) has important therapeutic implications. The manipulation of iron metabolism and the induction of ferroptosis are emerging as potential strategies in cancer therapy[58]. For instance, targeting the pathways associated with metallothioneins might sensitize tumor cells to treatments like chemotherapy or radiation, which rely on the production of Reactive Oxygen Species (ROS) ([59]).

### SCTP-CRC recognises shared and specific early signal in the tumor microenvironment of liver metastases from CRC

Furthermore, we applied SCTP-CRC to ST data 4 in Table 3, collected from two CRC liver metastasis (LM) samples, labeled as L1 and L2, with the purpose of investigating the applicability of SCTP-CRC for tumor microenvironment prediction in the context of metastases. According to the predicted tumor likelihood for sample L1 and L2 respectively (left panel of Figure 6A and B), SCTP-CRC demonstrated a substantial concordance with the diagnostic outcomes derived from pathological segmentation, which was evidenced by a considerable overlap of the predicted and actual tumor regions. To explore the origination and developmental trajectory of the metastasis, we employed Monocle3 ([60]) for a trajectory analysis on the predicted tumor spots (Methods), which chronologically mapped the estimated pseudotime to predicted tumor regions on the histological images (right panel of Figure 6A and B). Based on the estimated pseudo time, we defined different regions (i.e. Root, Invasion, Front and Normal as described in Methods). This time mapping evidently shows the trajectory from the root (core region) of the metastasis and outlined the invasion route. Intriguingly, we identified a distinct subset of tumor spots as a separate cluster in the trajectory analysis (marked in dark red), which appeared not to be part of the main invasion trajectory. We inferred that this cluster likely represent the invasion front or the interface within the tumor microenvironment, by correlating these spots with location distribution on the histological images.

**Figure 6:**
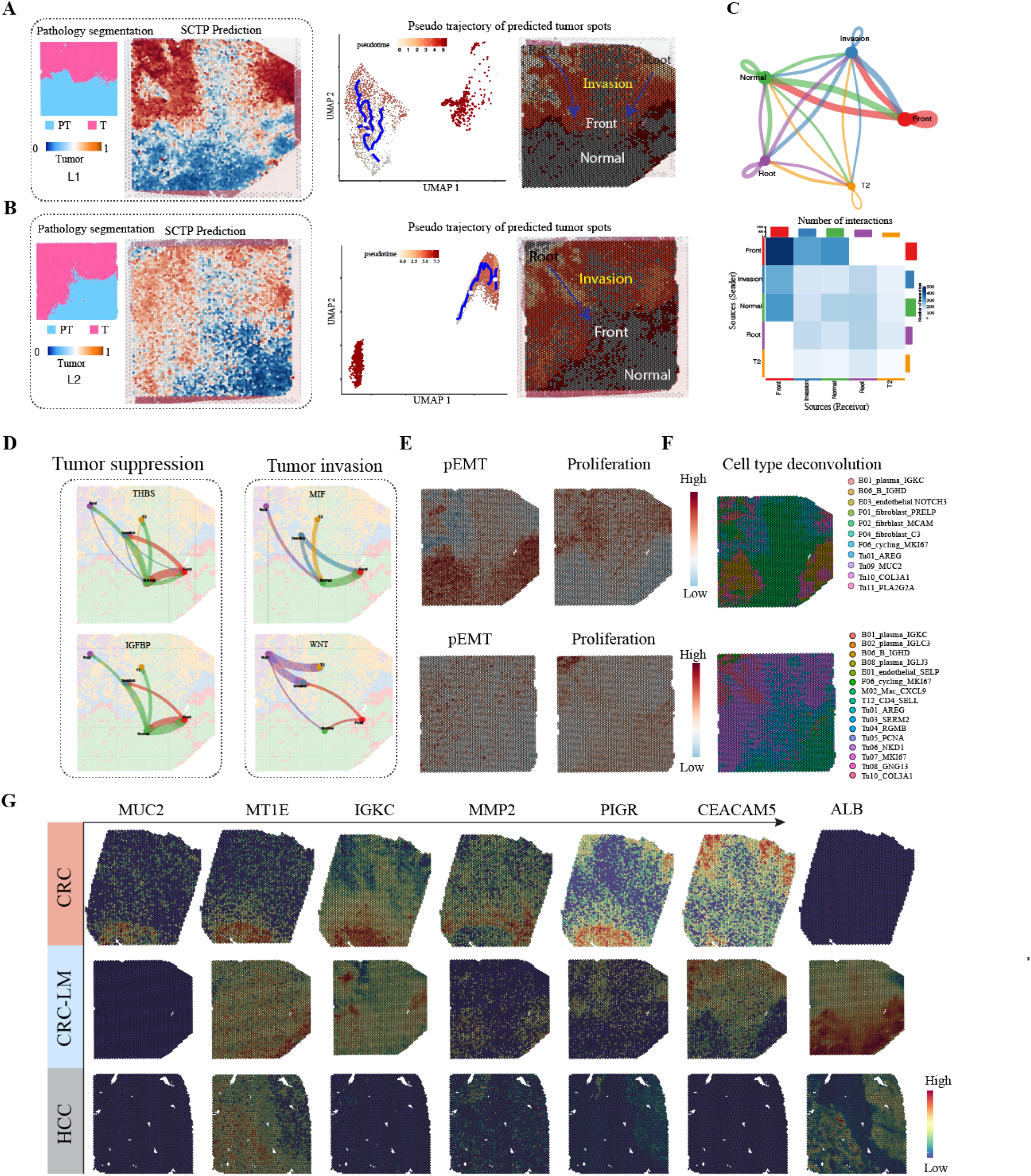
SCTP-CRC recognises shared and specific early signal in the tumor microenvironment of liver metastases by transformative prediction. **A, B** Left: Predicted tumor and peri-tumor areas for two spatial datasets L1 and L2 respectively using SCTP-CRC model. Right: Trajectory analysis on predicted tumor spots. The estimated pseudotime was then mapped to the histology images. **C** Inferred Cell-cell communication network visualized by circle plot and heatmap which shows the numbers of interaction between any two cell groups defined on pseudotimes from **A. D** Significant signaling pathways of individual L-R pairs from **C. E** Expression level of genes in pEMT and Proliferation pathways. **F** Cell composition which shows the most dominant cell cluster. **G** Expression level of genes MUC2, MT1E, IGKC, MMP2, PIGR, CEACAM5 and ALB in ST samples Q4, L1 and a primary HCC sample.

Next, we investigated the ligand-receptor pairs between five distinct subregions in sample L1 using CellChat v2 ([61]) (Methods). There are strong interactions identified between Front (interface) and Invasion / Normal areas, which suggests active cell-to-cell communication in these regions (Figure 6C). According to the remarkable signaling passes (the left part of Figure 6D), there was signal transmission among cells from Front to Invasion area, encompassing Thrombospondin (THBS) and Immunoglobulin Binding Protein (IGFBP). THBS is implicated in the tumor suppression pathway, and one of its isforms, THBS1, has recently been shown that it can produce tumor-infiltrating monocyte-like cells with contribution to immunosuppression and metastasis in colorectal cancer [62]. IGFBP plays a role in immune protection, and some proteins, including IGFBP-3 and IGFBP-rP1, have shown to be tumor suppressor in prostate cancer ([63, 64]), where our finding might confirm its similar role in liver metastasis of CRC. Conversely, signals indicative of tumor invasion (Figure 6D, and Supplementary Figure 4A) were observable in the opposite direction, including WNT ([65]), NOTCH and VEGF pathways with extensive research in the context of tumor invasion and metastasis [66]. WNT is a crucial signaling pathway during tumor invasion, and promotes migration and epithelial-mesenchymal transition (EMT) and facilitates proliferation and apoptotic resistance of CRC cells [67]. In colorectal cancer, antiangiogenic therapies, particularly those targeting vascular endothelial growth factor (VEGF) to inhibit new blood vessel formation in tumors, are generally used [68]. Besides, Notch signaling plays a role in cancer metastasis [69].

### SCTP-CRC detects key marker genes and independently assesses their association with colorectal cancer development and metastases

According to the above SCTP-CRC predictions, we identified DEGs with particular expression in tumor regions of the tumor microenvironment (Suuplementary Data 4), and the signature scores also showed different pathway activity in the tumor compared to other regions, such as cellular proliferation and pEMT, showing agreement with SCTP predictions (Figure 6E). Especially, the cells of Tu09 MUC2 were concentrated in the Front (Interface) region based on cell deconvolution (Figure 6F), so that, we found again the existence of Tumor MUC2+ in liver metastases of CRC. This led to our speculation that MUC2 and other above key marker genes might be specifically expressed in early stage of colorectal cancer and its liver metastases (CRC-LM), thus we compared the expression levels of key genes (e.g. MUC2, MT1E, IGKC, MMP2, PIGR) and liver-specific gene ALB, between CRC, CRC-LM and primary hepatocellular carcinoma (HCC) samples from ST data 5 in Table 3 (Figure 6G). In agreement with above findings, MUC2 is significantly expressed in Tumor MUC2+ in CRC sample Q4 and no expression was detected in liver metastases L1 and HCC samples. For other new marker genes, MT1E is highly expressed in Tumor MUC2+ in Q4, tumor margin in L1 and transient state in HCC ([70] Fig. 3C). MT1E could induce apoptosis and suppress the metastasis of HCC cells ([71]), suggesting Tumor MUC2+ is actually in early stage in colorectal cancer and show metastasis inhibition. IGKC and MMP2 are highly active at the Interface between the Tumor MUC2 and Normal regions, and the two genes also have signal detected in Normal and Interface area of liver metastases, but no in primary HCC. Meanwhile, PIGR and CEACAM5 are highly expressed in the Tumour regions of primary CRC and its liver metastases, while their signals are lower in primary HCC. PIGR is markedly overexpressed in the peritumoral region, which can mediate the transport of polymeric immunoglobulin across mucosal epithelial cells and is a key component of the mucosal immune system that bridges innate and adaptive immune defense [72]. Finally, the ALB gene is an important gene in the context of liver metastasis, serum albumin has utility as a prognostic indicator of cancer survival [73].

Finally, we conducted an external validation of SCTP-CRC tumor region prediction and above key genes on separate ST datasets of primary CRC ([74]) (ST data 3 in Table 3). Again, a notable concordance was observed between the tumor areas predicted by SCTP-CRC and those annotated in original study as shown in Supplementary Figure 4B. Remarkably, the expression levels of key genes such as MUC2, MT1E, IGKC, and PIGR aligned well with our above findings, reinforcing the validity of SCTP-CRC predictions and substantiating the accuracy of SCTP-CRC in identifying tumor-relevant regions and marker genes. The differential expression of these key genes underscores the heterogeneous nature of tumor cells and their interplays with surrounding tissues. Meanwhile, the expression of established CRC biomarkers, such as VEGFA and IGFBP2, was observed to be significantly increased in areas predicted to be tumors (Supplementary Figure 4C). However, some other biomarkers, including KRAS and TP53, did not show clear regional distribution (Supplementary Figure 4D), suggesting that they have limited efficiency in identifying tumor location in spatial dimension. These cellular gene expression profiles not only highlight the complexity inherent in the microenvironment of CRC and its liver metastases, but also shed light on the molecular dynamics that drive tumor growth and the corresponding host response. These results suggest that MUC2, MT1E and IGKC should play an important role in the early establishment of the microenvironment construction during CRC development or metastases.

## Discussion and conclusion

There are an increasing number of studies reporting significant scientific discoveries about the functions of various cell types in cancer through scRNA-seq or spatial transcriptomics. However, it is still challenging to predict the phenotype (e.g., malignancy) of individual cells or tumor patches, resulting in a notable hurdle for the accurate study of the tumor microenvironment. Accurate tumor probability predictions for individual cells or patches could help understand cellular heterogeneity within the tumor microenvironment and uncover molecular heterogeneity for disease diagnosis. In fact, the macro-phenotype information (such as ’Tumor’ versus ‘Normal’) can be collected from bulk data. However, how this key information can be translated to interpret the micro-phenotypes of individual cells and spatial spots remains an open question.

In this work, we introduce SCTP, a new machine learning approach that brings together phenotype information from bulk data, cell composition information from scRNA-seq data and spatial spots information from ST data. When ST data is not available, SCTP becomes a single modality model for single-cell analysis, where it represents an evolution that builds on and extends the capabilities of existing methods such as Scissor ([20]) and scAB ([21]). Deviating from these methods, SCTP uniquely integrates graph-based GAT machine learning techniques to account for cell connectivity, enabling a more nuanced understanding of cellular interactions. Further more, comparing to other graph-based algorithms like GCN, the attention mechanism of GAT allows for allocating more weights to important cells and spots, therefore provides more accurate prediction. This graph integration, particularly when supplemented with ST data, significantly increases the effectiveness of SCTP in identifying phenotypic cell subpopulations, surpassing traditional methods in both scale and precision. Moreover, SCTP is computationally more efficient compared to transfer learning based method such as SpaRx ([23]) that learns the predictive model from bulk RNA-seq data and transfer it to single cell spatial transcriptomic data. Since SCTP takes the correlation matrix between bulk data and single cell/spatial data as the input, the number of features in our model corresponds to the bulk sample size, which is usually much lower comparing to the number of genes sequenced in RNA-seq data. In their recent work, [75] present Cancer-Finder, a novel framework designed to identify tumor regions with high accuracy. This approach leverages a pre-trained model applied to annotated spatial transcriptomics (ST) data. Despite the innovation, a significant limitation arises from the scarcity of readily available annotated ST datasets, coupled with the framework’s reliance on binary predictions, which may oversimplify the complexity of tumor heterogeneity.

The decision to use specific CRC datasets for SCTP-CRC construction was driven by several factors. First, the large number of cells provides a solid basis for analyzing different cell types and their interactions. This large cell number increases the statistical power of our analyzes and enables more reliable and generalizable results. Second, the rich diversity of cell types is particularly advantageous, providing a comprehensive overview of the cellular landscape that includes both immune and non-immune cells. This diversity is crucial for SCTP-CRC, which aims to explore complex cellular interactions and the molecular basis of various cellular processes. Third, both the single cell and spatial data in the SCTP-CRC are sourced from the same study ([45]) that was most recently conducted, which provides data with higher precision and coverage. Moreover, samples from the same cohort guaranties the homogeneity across different data sources.

Of note, in SCTP, the parameter *α* plays a pivotal role in the predictive model, acting as a balancing factor between scRNA-seq data and ST data. Its value, lying within the range of 0 to 1, modulates the relative influence of each data type on the final prediction. Specifically, as *α* approaches 1, the contribution of scRNA-seq data becomes more critical in the determination of predictive outcome. Conversely, a value nearing 0 indicates a greater dependence on ST data. Importantly, *α* can be customized to suit the specific aims of a study, allowing researchers to tailor the model’s sensitivity to the distinct properties of each data type. Another parameter *c*, determining the number of meta-cells to include in the training data, is adjustable as well. Although a large value of *c* might contain more information, it will also increase the computational burden and weaken the signal from spatial transcriptomic data. In practice, we advise using a comparative number of cells to the total number of spots gathered in the ST data.

SCTP is designed to integrate both single-cell and spatial transcriptomic data, yet is flexible enough to work with only one of these data sources. This adaptability is demonstrated in our single-modality method comparison, where we provide an example of SCTP using only single-cell data and outperforming conventional Scissor and scAB methods. In other scenarios using only spatial transcriptomic data, we found the efficiency of tumor area prediction is limited because the predicted value is close to binary and less informative due to the lack of higher resolution data on cells. Although single-modality spatial transcriptomics can comprehensively identify tumor regions, the finer resolution provided by scRNA-seq data is critical to obtain more precise and accurate phenotypic predictions from SCTP.

The key of SCTP is to merge the RNA-seq and scRNA-seq data into ST data, so that, it should provide a high resolution of tumor microenvironment in a phenotype viewpoint. As a representative study on colorectal cancer, we showcase the wide-ranging use of SCTP. SCTP first shows its exceptional ability to classify cells or spots precisely as malignant or non-malignant. On cell level, SCTP precision is evidenced in its successful prediction of cells with different colorectal cancer subtypes (i.e. CSM1, CSM2, and CSM3) and tumor marker genes, along with a high degree of correlation between SCTP’s predictions and pathological diagnoses. Especially, SCTP identified a hidden area in a primary colorectal sample, which has not been recognised by conventional pathology segmentation. According to DEGs associated to this area, it would be an area in early tumor state / stage. Several key DEGs were also validated their expressions and spatial distributions in independent ST data of CRC. Besides, the liver emerges as the primary site for metastasis in cases of colorectal cancer. Approximately 50% of patients either exhibit liver metastases concomitantly at the time of diagnosis (termed synchronous metastasis) or manifest liver metastases in a delayed manner throughout the disease trajectory. Dependent on SCTP prediciton, pseudotime examination of predicted tumor regions in liver metastases of CRC provided a clear representation of the invasion route, which was further elucidated through the ligand-receptor interactions across different invasion stages. Most above key DEGs were also seen between predicted tumor and peri-tumor areas, and were corroborated by external studies. These series of results indicate that SCTP should be a valuable machine learning tool to detect new targets for early diagnosis of CRC or other complex diseases ([76]). Of note, the cells and spots analyzed would ideally come from samples exhibiting the same phenotype as bulk data, such as colorectal cancer in our study. However, this is not a prerequisite for the application of SCTP. For example, this flexibility of SCTP-CRC is exemplified that phenotype of CRC was initially characterized and further successfully transferred to make predictions on liver metastases of CRC.

In summary, SCTP is a novel tool for phenotype prediction using single-cell and spatial gene expression data. It fills a critical gap in automated analysis by accurately estimating macro-phenotype (e.g. malignancy) at the molecular and cellular levels, which is critical for studying the tumor microenvironment. SCTP-CRC has shown great success in studying colorectal cancer and its liver metastases, providing a scheme for the discovery ”key DEGs *→* new subtypes of CRC associated cells *→* cellular spatial distribution”. Furthermore, the accurate malignant and non-malignant prediction on spatial samples offers the possibility of locating the interface where tumours contact the microenvironment. This information is crucial for comprehending how tumours spread into neighbouring tissues ([77]), and how to accurately define this interface from the prediction of SCTP remains a topic worth further investigation. Beyond its application in tumor prediction, SCTP holds potential for studying drug resistance and other complex biological scenarios, paving a new path for personalized medicine in clinical applications.

## Methods and materials

### SCTP model

The Single-Cell and Tissue Phenotype prediction (SCTP) framework, through its multi-task fusion learning model, aims to simultaneously predict phenotypes of specific cells and spatial spots by guidance from bulk sample and data. SCTP consists of data preparation, multi-scale network construction, multi-task learning with latent space fusion, and phenotype prediction. Especially, the learning network structure of SCTP incorporates a parallel within-modality Graph Attention Network (GAT) alongside a parallel Multi-Layer Perceptron (MLP), where a cross-modality loss for coefficient distribution fusion is utilized.

### Data preparation

There are three data sources for SCTP (Figure. 1A), a single-cell expression matrix *M*_*sc*_ of *k* cells, a spatial expression matrix *M*_*st*_ with a tissue histology image cut in *l* spots, and a bulk expression matrix with *n* samples with phenotypes *Y* = (*y*_1_, …, *y*_*n*_) of interest.

### Multi-scale network construction

On one hand, the network for cell-cell similarities **G**_*sc*_ was calculated by the shared nearest neighbor (SNN) algorithm ([78]), which determines the similarity between cells by taking into account both the degree of direct proximity between cells and the degree to which they share neighbors. The matrix **X**_*sc*_ *∈ R*^*k×n*^, representing the correlation between individual cells and the bulk samples, was accomplished by computing Pearson’s correlation between bulk RNA-seq data and scRNA-seq data. On the other hand, the networks for spot-spot similarities **G**_*st*_ were created by SpaGCN ([79]), which integrates spatial location with histological information to guarantee a thorough evaluation of the areas being studied. The correlation matrix **X**_*st*_ *∈ R*^*l×n*^, representing the correlation between individual spots and the bulk samples, was accomplished by computing Pearson’s correlation between bulk RNA-seq data and ST data (Figure. 1B). The correlation matrix **X**_*sc*_ and **X**_*st*_ are assigned to the created graphs **G**_*sc*_ and **G**_*st*_ as features of each node.

### Multi-task learning

To efficiently process and synthesize data from above two graphs, SCTP combines a parallel Graph Attention Network (GAT) ([80]) with a Multi-Layer Perceptron (MLP) ([81]), i.e. the parallel GAT blocks are followed by two MLP embedding layers. The parallel attention operation takes two graphs **G**_*sc*_ and **G**_*st*_, together with the corresponding node features **X**_*sc*_ and **X**_*st*_ as inputs. Taking the single cell data as example, where **X**_*sc−i*_ denote the input feature of cell *i*, i.e. the correlation between cell *i* and the bulk samples based on gene expressions. The self-attention mechanism computes the attention coefficients *a*_*ij*_ for each pair of cells *i* and *j* by:

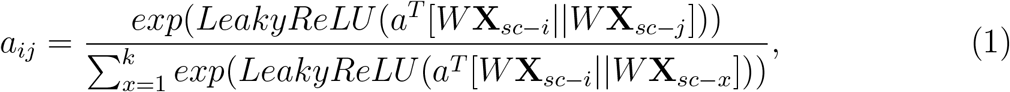

where *a ∈ R*^2*k*^ is a learnable weight vector, 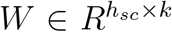 is a learnable weight matrix with *h*_*sc*_ the hidden size, || is the concatenation operation and *LeakyReLU* is the leaky rectified linear unit activation function. These attention coefficients are used to weight the messages of a cell’s neighbors, which are the neighbor’s features multiplied by the same weight matrix *W*. The output feature of cell *i* is therefore obtained by:

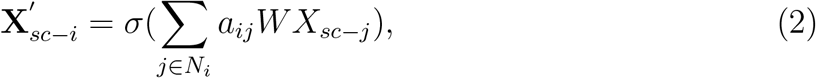

where *N*_*i*_ is the set of the neighbours of cell *i* and *σ* is a nonlinear activation function. When employing the multi-head attention mechanism, suppose *M* independent attention mechanisms execute the transformation of Equation (2), their features are concatenated and result in the following output feature representation:

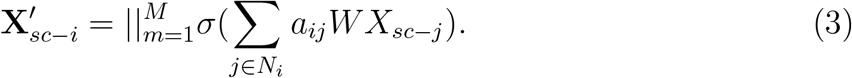

The attention mechanism allows to determine the importance of each cell, assigning more weight to the pertinent ones while reducing the emphasis on less relevant cells. As a result, this empowers the model to improve prediction accuracy by concentrating its attention on the most critical information. Afterward, we derive the cell embedding 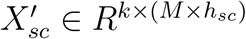. This intermediate embedding is subsequently refined in the following MLP phase to generate the final output. An MLP is a fully connected multi-layer neural network that learns a function *f* (*·*) : *R*^*d*^ *→ R*^*o*^ through training on input data, where *d* is the number of dimensions for input and *o* is the number of dimensions for output. In our case, this architecture consists of an input layer utilizing the embedding 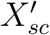 derived from GAT with *d* = *M × h*_*sc*_, followed by a hidden layer of size *h*, and an output layer containing a single node (*o* = 1). The final output is a vector of length *k*, denoted as *β*_*sc*_, representing the aggregated information pertaining to each cell.

The approach for processing spatial transcriptomics (ST) data follows a similar procedure. Initially, we employ GAT with a hidden size of *h*_*st*_ to obtain the intermediate output features for spots, resulting in a feature matrix of size *l ×* (*M × h*_*st*_). Subsequently, an MLP is applied to yield the final embedding for all the spots, represented as *β*_*st*_. The coefficients *β*_*sc*_ and *β*_*st*_ serve as latent space indicators of the phenotypic association of each cell or spot and capture significant phenotypic data that correlates with bulk sample. A higher significant phenotypic relevance is indicated by higher coefficients.

### Latent space fusion

With estimated coefficients *β*_*sc*_ and *β*_*st*_, phenotype prediction is achieved by a linear combination of these indicators, which represent synthesized information from scRNA-seq and ST data. This can be considered as a information fusion of two data sources, and provides the probability of the phenotype *Y* = 1 (for example, in a binary classification task where phenotype is coded as 0 and 1):

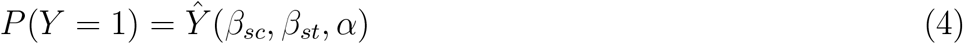

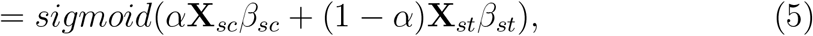

The *sigmoid* function is employed to adjust the predictive values to a range of 0 to 1, which describes the likelihood that the prediction will fall into class 1. The parameter *α* is a penalty parameter to balance the attention on scRNA-seq or ST data.

To estimate the coefficients *β*_*sc*_ and *β*_*st*_, cross-entropy is employed as a loss function in the model optimization ([82]). In the case of binary classification task, the cross-entropy between observed phenotype *Y* and its prediction can be written as:

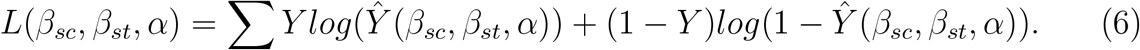

Given a fixed *α*, two coefficient vector *β*_*sc*_(*α*) and *β*_*st*_(*α*) are estimated by optimizing this loss function.

Of note, to construct a more accurate and informative predictor, *α* was chosen such that the fusion of *β*_*sc*_(*α*) and *β*_*st*_(*α*) minimizes the distance between the distribution of *β*_*sc*_(*α*) and *β*_*st*_(*α*), reflecting similar phenotype distribution in fusion latent space. Suppose *β*_*sc*_(*α*) and *β*_*st*_(*α*) follow distribution 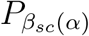 and 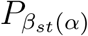 in the same latent space *𝔛*, the distance between 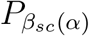 and 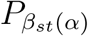 can be measured by Kullback-Leibler (KL)*−*divergence ([83]), such as:

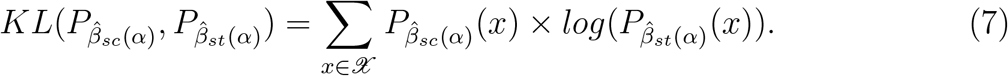

The parameter *α* is then optimized to minimize this distribution distance:

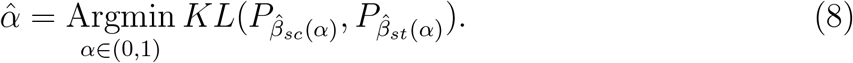

In practice, we search through a grid between 0 and 1 to find the optimal value of *α*.

### Training procedure for multi-modal prediction

For SCTP implemented in this work, the GAT comprises a multi-head GAT architecture with *M* = 3 different heads, where the *n* features, corresponding to the expression correlation with the *n* bulk samples, are embedded into hidden vectors with *h*_*sc*_ = *h*_*st*_ = 96. MLP consists of two fully-connected layers, with a Gaussian Error Linear Unit (GELU) activation function between them. The first layers contain *h* = 48 hidden nodes, and the outputs of the second layers are one-dimensional vectors *β*_*sc*_ and *β*_*st*_. At the end, a linear layer and a sigmoid layer, denoted in Equation (4), can be used for classification task. The model undergoes training over 1000 epochs using an Adam optimizer with a starting learning rate of 0.002, which decays at a factor of 0.99 each epoch, and employees cross-entropy loss as the loss function.

The most important output from the above training procedure is two optimized coefficients 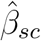 and 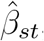, alongside 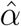 which is chosen following Equation (8). These coefficients and parameters are essential for phenotype prediction in new scRNA-seq and ST samples. Two matrices, the gene expression matrix from single cells and spatial spots, denoted as *M*_*sc*_ and *M*_*st*_, are also required for prediction. Denote the new gene expression of a cell or spot as *g*, the correlation vector between *g* and *M*_*sc*_ is calculated as *ρ*_*sc*_ = *corr*(*g, M*_*sc*_), and the correlation between *g* and *M*_*st*_ is calculated as *ρ*_*st*_ = *corr*(*g, M*_*st*_). The correlation vectors *ρ*_*sc*_ and *ρ*_*st*_ are then inputted into the SCTP model with estimated parameters (or as network weights) of 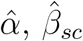 and 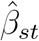:

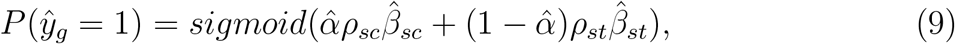

In the case where phenotype ’tumor’ is coded as 1 and ’normal’ is coded as 0, the predicted value can be considered as the tumor likelihood. Prediction closing to 1 indicates that the cell is likely to be tumor cells or the spot comprises tumor cells in that region.

### Single-cell subpopulation identification by phenotypic coefficients

#### Individual cell identification from single cell RNA sequencing

The coefficient 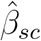 is crucial for distinguishing cell subpopulations associated with the specific phenotype of interest, based on scRNA-seq data. Cells with 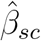 values exceeding the 90_*th*_ percentile are defined as phenotype-relevant, indicating a strong association. Conversely, cells with values below this threshold are categorized as background cells, signifying a lesser or negligible association.

#### Individual spot identification from spatial transcriptomics

Unlike scRNA-seq data, the histology image coming with the ST data offers enhanced visualization capabilities and allows for direct interpretation from 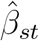. Positive values of the estimated coefficient 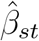 are indicative of a positive correlation between spatial locations and the phenotype being investigated. Conversely, negative values of 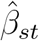 suggest a negative correlation. These coefficients thus serve as indicators of the direction and strength of the association between spatial spots and the phenotype under study. In scenarios where the classification differentiates between tumor and normal samples, spots with positive 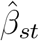 values are potentially indicative of tumor regions. Conversely, areas exhibiting negative values of 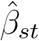 are likely to correspond to normal tissue. This distinction provides a valuable tool for spatially mapping tumor and normal tissues based on their association with the phenotype, as indicated by the coefficient 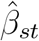.

### Meta-cell representation and selection

In scRNA-seq data, there are an abundance of cells usually. Such high volumes pose challenges for data analysis and model training, particularly when attempting to integrate these cells with a much smaller number of spots. As recently reported in [84], meta-cell based on a downsampling technique could be effectively incorporated to address data imbalance and expedite the analysis process. Specifically, the set of meta-cells can still exhibit the highest connectivity, as these are most representative and informative for training model and downstream single-cell analysis. By focusing on cells with the most significant interactions or relationships within others, meta-cells are selected for SCTP, which can help to effectively train model with fewer cells without compromising the integrity and depth of study.

The meta-cell selection has the following steps:

1. **Network construction using Shared Nearest Neighbors:** Let *G* = (*V, E*) be a graph where *V* represents the set of nodes (cells) and E represents the set of edges between these nodes. The network is constructed such that an edge *e*_*ij*_ *∈ E* exists between nodes *v*_*i*_ and *v*_*j*_ if and only if they are shared nearest neighbours.
2. **Degree Calculation**: For each node *v*_*i*_, its degree *d*(*v*_*i*_) is defined as the number of nodes that are connected to *v*_*i*_ by an edge in *V*.
3. **Cell Clustering and Selection**: If cell type annotations are available, then each cell type forms a distinct cluster. Otherwise, the clusters are defined by the hierarchical agglomerative clustering Louvain algorithm ([85]). Assume the cells are clustered into *s* different clusters, *C*_1_, *C*_2_, …, *C*_*s*_. For each cluster *C*_*i*_, the cells that have the top *c* highest degrees within that cluster, are denoted as *S*(*C*_*i,c*_).
4. **Final meta-cell selection** The final selection of meta-cells *S* is the union of the above selected cells from each cluster, i.e., *S* = *∪*_*i*=1,…,*s*_*S*(*C*_*i,c*_)

The parameter *c* is adjustable to ensure that the total number of selected cells (from scRNA-seq data) approximates the number of spots (from ST data).

### Single modality model of SCTP using scRNA-seq and bulk RNA-seq data

SCTP model is proposed to integrate bulk data with single-cell data and spatial data. Notably, the absence of one modality (either single-cell or spatial data) does not impede the functionality of SCTP. Consistent to previous methods (e.g. Scissor or scAB), we consider a scenario where only scRNA-seq data is available, analogous to cases with exclusively spatial gene expression data. When equipped with bulk RNA-seq data in addition to scRNA-seq, a graph **G**_*sc*_ along with its node features **X**_*sc*_ is constructed. These elements are subsequently processed through a GAT, facilitating the generation of embeddings for single cells based on their correlation with bulk data. Subsequent to this, a MLP is employed to estimate the coefficient *β*_*sc*_. The SCTP prediction is then generated as follows:

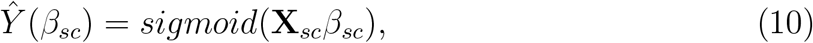

where *β*_*sc*_ is estimated by minimizing the following loss function:

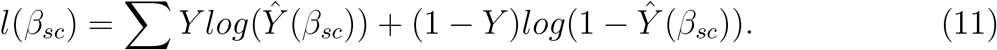

### Datasets

#### Bulk RNA-seq datasets

On one hand, bulk RNA-seq dataset for immunotherapy, PRJEB23709, was provided by [86], and two bulk RNA-seq datasets on immunotherapy, MGSP (Melanoma Genome Sequencing Project, [87]) and GSE91061 ([33]), are served as validation set. On the other hand, bulk RNA-seq data for colorectal cancer was used as the GSE44076 ([27]) dataset, providing gene expression profiles of paired normal adjacent mucosa and tumor samples from 98 CRC individuals. Three datasets, GSE18105 ([30]), GSE21510 ([31]), GSE23878 ([32]) that collected samples with primary colorectal cancer and normal are sourced as validation set. Information on these data can be found in Table 1.

#### scRNA-seq datasets

The scRNA-seq dataset for melanoma patients was obtained from GSE115978 (31 melanoma tumors), which contains 6879 cell of nine cell types: B cells, cancer associated fibroblasts (CAFs), endothelial cells, macrophages, malignant cells, Natural Killer (NK) cells, CD4+ T cells, CD8+ T cells, and T cells ([29]). For a deep case investigation, three CRC-associated single-cell datasets were included. One is derived from GSE146771 ([25]), collecting from tumors, adjacent normal tissues, and blood of 18 treatment-naive CRC patients. This dataset includes 43817 cells previously clustered in five cell types (B cells, CD4 T cells, CD8 T cells, ILCs and myeloid cells). The second one is from GSE132465 ([26]), which collected a total of 63689 cells from 23 Korean colorectal cancer patients and 10 matched normal mucosa. Cell types including Epithelial cells, T cells, B cells, myeloid cells, stromal cells, and mast cells can be found in this dataset. The last one is sourced from the research conducted by [45], encompassing a comprehensive collection of 41892 CD45*−* nonimmune cells and 196473 CD45+ immune cells derived from 27 samples with primary colorectal cancer, adjacent normal colorectal mucosa, liver metastasis, adjacent normal liver tissue, and peripheral blood. Information on these data can be found in Table 2.

#### Spatial transcriptomics datasets

For the application of SCTP in deep case study of CRC, ST data were collected from two recent studies. The first dataset was obtained from the study conducted by [34], which contributed four distinct ST data combining histological information (labeled P1-P4). The second dataset was originated from the investigation by [45], comprising four ST data of primary CRC (labeled Q1-Q4) and two ST data of liver metastatic tumors of CRC (labeled L1-L2). For each sample, the tumor area (T) and adjacent paratumor area (PT) were identified using hematoxylin and eosin (H&E) staining and gene expression features as pathological diagnose according to their original publications.

Besides, we employed ST data from [74] as an external validation set for CRC case, which pertains to primary CRC. From this dataset, we selected two samples (labeled as S1 and S2) that exhibited clear tumor margins as reported in the original study. The SCTP-CRC model was applied to predict tumor likelihood of each spot within these samples. These predictions were then compared with the pathologist annotations provided in the original study. Additionally, to assess the specificity of the marker genes identified in CRC and its liver metastases, we incorporated ST data from a study on primary liver cancer ([70]), and selected one sample from this dataset, particularly a leading-edge section where tumor regions and normal tissues are juxtaposed. Information on these data can be found in Table 3.

#### TCGA validation dataset

To provide gene expression data of a large cohort for marker gene validation, we also applied the RNA-seq data from the TCGA-COAD (The Cancer Genome Atlas-Colon Adenocarcinoma), which included 437 samples that were explicitly annotated with a defined cancer stage (e.g. Stage I - IV). The sample size of different stages are I (81), II (167), III (123) and IV (66) respectively.

### Data preprocessing

Bulk RNA-seq data were downloaded and normalized, and corresponding phenotype data was also collected.

The Seurat R package (version 5.0.0) was utilized to process scRNA-seq and ST data. Consistent with typical single-cell data analysis protocols, our initial step involved removing genes that demonstrated low expression levels across cells and also removing cells with low genes and UMIs. Subsequently, the remaining expression data was normalized utilizing the *NormalizeData* function with the default parameters. We then identify highly variable genes through the *FindVariableFeatures* function. Following this, a transformation of the data was conducted using the *ScaleData* function, and principal component analysis was applied to these scaled data using the *RunPCA* function. On the first ten principal components, the next step involved calculating cell-cell similarity based on shared nearest neighbor graph by the *FindNeighbors* function, and the resulting cell-cell similarity matrix underwent a binarization process with diagonal elements as zero. Finally, we utilized the *RunUMAP* and *RunTSNE* functions to perform dimensionality reduction and to visualize cells in low dimensions.

For ST data, we first create a Seurat object for each sample within downloaded data, which contains the histology image of the sample, the coordinate of each spot and their gene expression. We then utilized the function *SCTransform* from R package Seurat for normalization on the expression matrix, with default parameters.

### SCTP-CRC model construction and evaluation

In the context of the SCTP application in colorectal cancer, the target variable, denoted as *Y*, is defined in binary terms: ”1” represents ”tumor,” while ”0” signifies ”normal”. The bulk data utilized in this study was sourced from GSE44076, comprising a total of 196 samples, evenly split between 98 tumor and 98 normal samples. Next, based on gene expressions, we predict the phenotype of non-immune cells (including tumour cells) and spots on ST data Q2 and Q4 from tissue specimens of CRC patients, as described in Equation 9. The predicted tumour likelihood are overlaid onto the tSNE plot for cells and the histology images for spots. Positive values are considered as malignant cells or tumor spots, while negative values are considered as non-malignant cells or peritumoral spots.

For cell predictions, we assess the accuracy of our predictions by comparing the predicted malignancy against the cell type annotations provided in the original study. Additionally, we compute the proportion of predicted malignant cells relative to non-malignant cells within each cell type. In the case of spatial data, we evaluate the SCTP-CRC predictions for each sample against the pathological segmentation of tumor and paratumor regions as documented in the original study.

## Downstream analysis for SCTP-CRC prediction

At the single-cell level, following malignancy predictions, standard downstream analyses typically include differential expression gene (DEG) analysis and pathway analysis. These steps involve identifying genes that are differentially expressed between malignant and non-malignant cells and then investigating the biological pathways associated with these genes. Such methodologies have been extensively covered in literature (e.g., [20], [21]). While acknowledging the significance of these established analysis, our study concentrates on the novel downstream analysis we performed on spatial spots. Once predictions on the spatial transcriptomic data are acquired, several subsequent analyses were conducted. Following the classification of spatial spots as tumor or normal using SCTP-CRC model, we embarked on a series of downstream analyses tailored to spatial transcriptomics.

### Differential expression gene analysis

We used the *FindMarkers* function in the Seurat package (version 5.0.0) to identify differentially expressed genes (DEGs) between tumour and normal spots, as predicted by SCTP, where the Wilcoxon rank sum test by default was used and two factors were adopted to identify DEGs: an absolute fold change in gene expression greater than one, and a statistical significance threshold based on a false discovery rate (FDR) less than 0.05. The relative expression levels of these genes were effectively visualised through the use of a volcano plot to illustrate the differential expression patterns.

### Signature score for cancer hallmark quantification and visualization

To elucidate the molecular functional diversity (e.g. cancer hallmarks) across various tumor regions, we utilized the *AddModuleScore* function from the Seurat package, which involved a pre-defined list of genes that represents a distinct functional category and cancer signatures.

### Cell-type deconvolution

At the current Visium ST resolution, each spot potentially contained approximately 8 to 20 cells ([88]). To estimate the proportions of cell types within each spot, we employed a deconvolution analysis using the Robust Cell Type Decomposition (RCTD) method as described by [44] and implemented in the Seurat package. RCTD is designed to leverage cell type profiles derived from scRNA-seq data to accurately decompose cell type mixtures in ST data, while also adjusting for variances attributable to differences in sequencing technologies. The analytical procedure commenced with the extraction of paired cells from scRNA-seq and ST datasets that were located within mutual neighborhoods. This was accomplished using the *FindTransferAnchors* function in Seurat, wherein the scRNA-seq data served as the reference and the ST data as the query. Subsequently, the identified ‘anchors’—correspondences between the two datasets—and the single-cell labels from the scRNA-seq data were utilized to perform data transfer using the *TransferData* function, executed with its default parameters. This method allowed for the projection of cell type labels from scRNA-seq data onto the ST spots, thereby facilitating a more precise understanding of cellular compositions within the spatially resolved tissue sections.

### Cell trajectory inference

Tumor cells exhibit varying gene expression profiles as they transition between different states, necessitating a process of transcriptional reconfiguration. Monocle ([60]), an advanced bioinformatics tool, has been designed to decipher the sequence of gene expression alterations that individual cells experience during dynamic biological processes. After capturing the overarching trajectory of gene expression changes, Monocle can accurately position each cell along this trajectory. To gain insights into the developmental trajectory and invasive characteristics of tumor cells during metastases, we employed the most recent version Monocle3 for the calculation of pseudotime corresponding to each predicted tumor spot with predicted tumor likelihood greater than 0.5. First, the function *learn graph* was used to construct the trajectory graph. Next, all analyzed cells were ordered according to its progress along this trajectory by the function *order cells*. The cells can be then colored by pseudotime which show their order. Finally, the estimated pseudotime for each spot can be extracted from the output of the function *learn graph*. Cells with infinite pseudotime indicate that they were not reachable from the root nodes. By mapping all the trajectory pseudotime onto the histology image, we could reconstruct the developmental sequence of tumor cells, tracing their progression from early to advanced stages. This approach can not only help visualize the invasive patterns of tumor cells, but also provides a temporal framework to understand the dynamic changes occurring within the tumor microenvironment.

### Cellular communication network analysis

To predict ligand-receptor pairings among distinct regions, we employed CellChat v2, a state-of-the-art tool developed by [61] for signaling inputs and outputs prediction. Initially, the predicted tumor region was segmented into various subregions based on their characteristics, e.g. the ‘Root’ (characterized by a high estimated pseudotime from trajectory analysis), ‘Invasion’ (comprising newly developed tumor cells), and ‘Front’ (encompassing the margin around the invasion region). More specifically, these groups are defined as follows:

1. **Root**: This region is defined by spots with an estimated pseudotime exceeding the 75th percentile (third quartile) of all pseudotime values, which represents spots indicative of the occurrence after long time within the tumor development.
2. **T2**: This region is characterized by spots that fall within the interquantile range of all pseudotime values. These spots are located within the tumor area, specifically between the root and the invading tumor cells.
3. **Invasion**: Comprising spots with estimated pseudotime below the 25th percentile of all pseudotime values. This area typifies the newly formed tumor at the margin, marked by expansion and invasiveness. It is a critical region in tumor progression, highlighting the initial phases of tumor expansion and spread.
4. **Front**: This region is identified by a distinct cluster in the time trajectory analysis, typically associated with an infinite pseudotime. This cluster represents the interface between tumor cells and normal cells, delineating a vital boundary in tumor development.
5. **Normal**: The region includes spots predicted to be peri-tumoral by SCTP-CRC with predicted tumor likelihood inferior or equal to 0.5. These spots, excluded from the trajectory analysis, represent areas unaffected by the tumor.

Subsequently, the *createCellChat* function from the package, as instructed by [61], was deployed to elucidate the ligand-receptor interactions across spots in these differentiated regions. Further, the *netVisual circle* function was used to generate circle plots, illustrating the strength of ligand-receptor interactions. And, the *netVisual heatmap* function was implemented to present the number of interaction pairs as a heatmap, facilitating an intuitive understanding of the interaction frequency and patterns. Finally, the *netVisual aggregate* function was applied to visually aggregate and highlight key signaling interactions, providing a comprehensive view of the cellular communication networks within the tumor’s microenvironment.

## Data availability

The following are the Gene Expression Omnibus accession numbers used: [27]: GSE44076; [33]: GSE91061; [25]: GSE146771; [26]: GSE132465; [29]: GSE115978; [30]: GSE18105; [31] GSE121510; [32]: GSE23878. The data PRJEB23709 and MGSP are available at https://github.com/donghaixiong/Immune cells analysis. TCGA-COAD data are available at the GDC portal (https://portal.gdc.cancer.gov). Primary liver cancer spatial transcriptomics data are available at http://lifeome.net/supp/livercancer-st/data.htm. The independent validation ST data of primary CRC is available at https://zenodo.org/records/7553463.

## Code availability

The code will be made available upon publication.

## Acknowledgement

This study was supported by the National Key R&D Program of China (No. 2022YFF1202100, 2023YFF1204700), the National Natural Science Foundation of China (No. 12371485, 11871456).

