## Supplementary Figure for "Detecting phenotype-specific tumor microenvironment by merging bulk and single cell expression data to spatial transcriptomics"

March 8, 2024

This supplementary material provides additional figures for the paper: "Detecting phenotype-specific tumor microenvironment by merging bulk and single cell expression data to spatial transcriptomics".

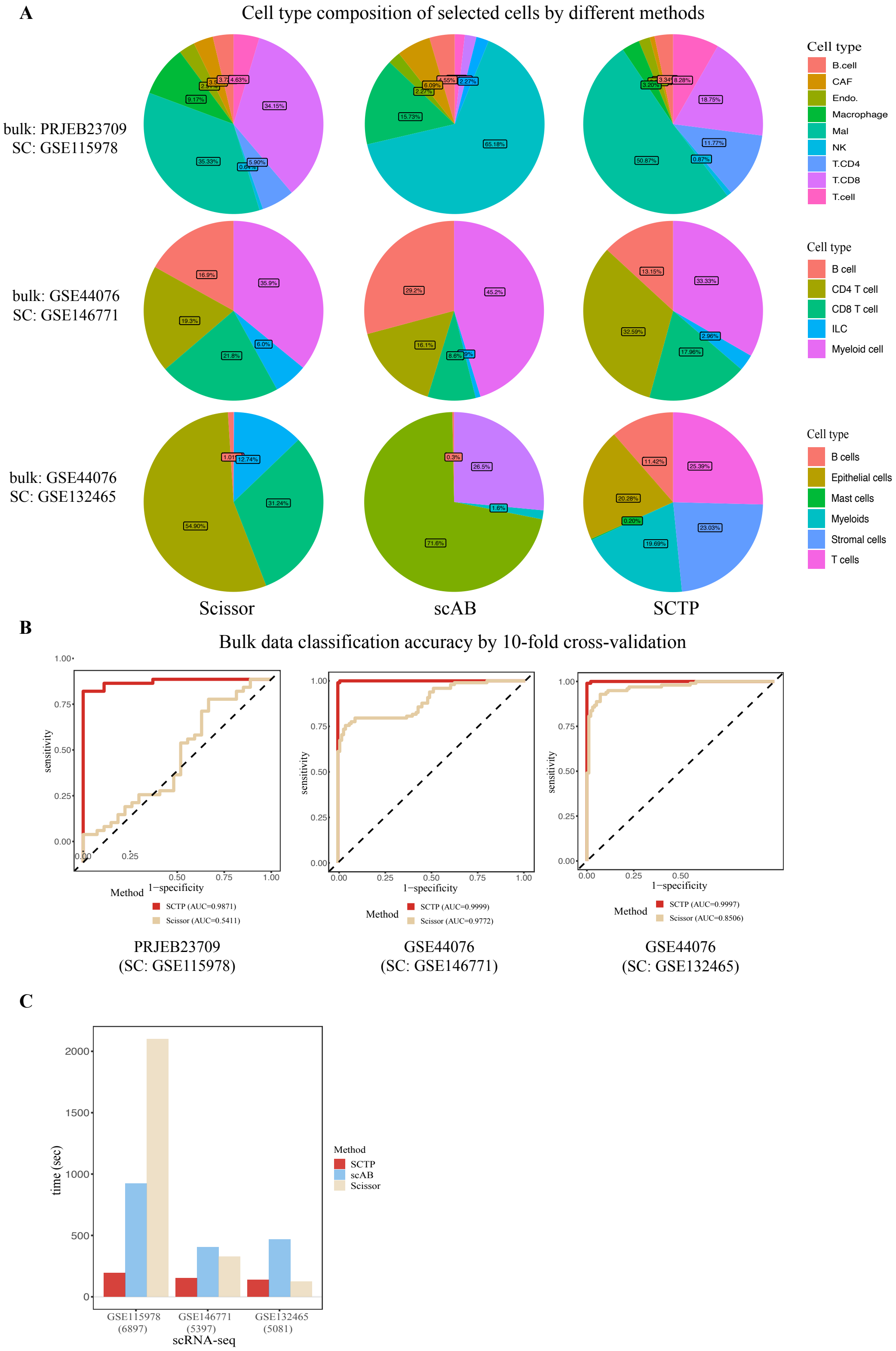


Figure 1: SCTP utilizing single modality in the form of single-cell RNA-seq data. **A**: Composition of the Selected Cell Populations. **B**: Accuracy of Bulk Data Classification Utilizing 10-Fold Cross-Validation. **C**: Computational expenses for SCTP, scAB, and Scissor. The number of single cells in different datasets are indicated in the parenthesis.

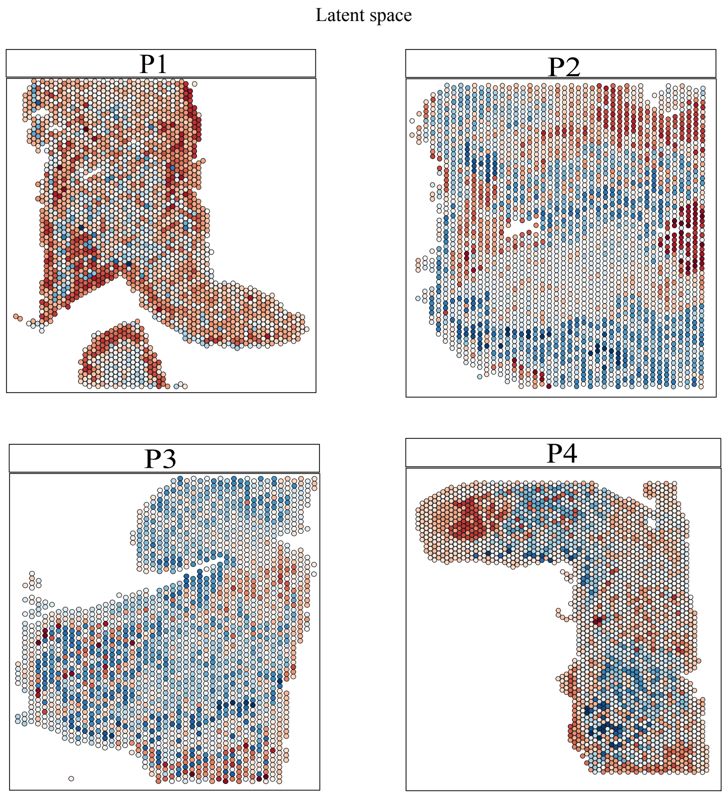


Figure 2: SCTP utilizing single modality in the form of spatial transcriptomic data. The estimated coefficients $\beta_{st}$ on the latent space in four datasets P1-P4.

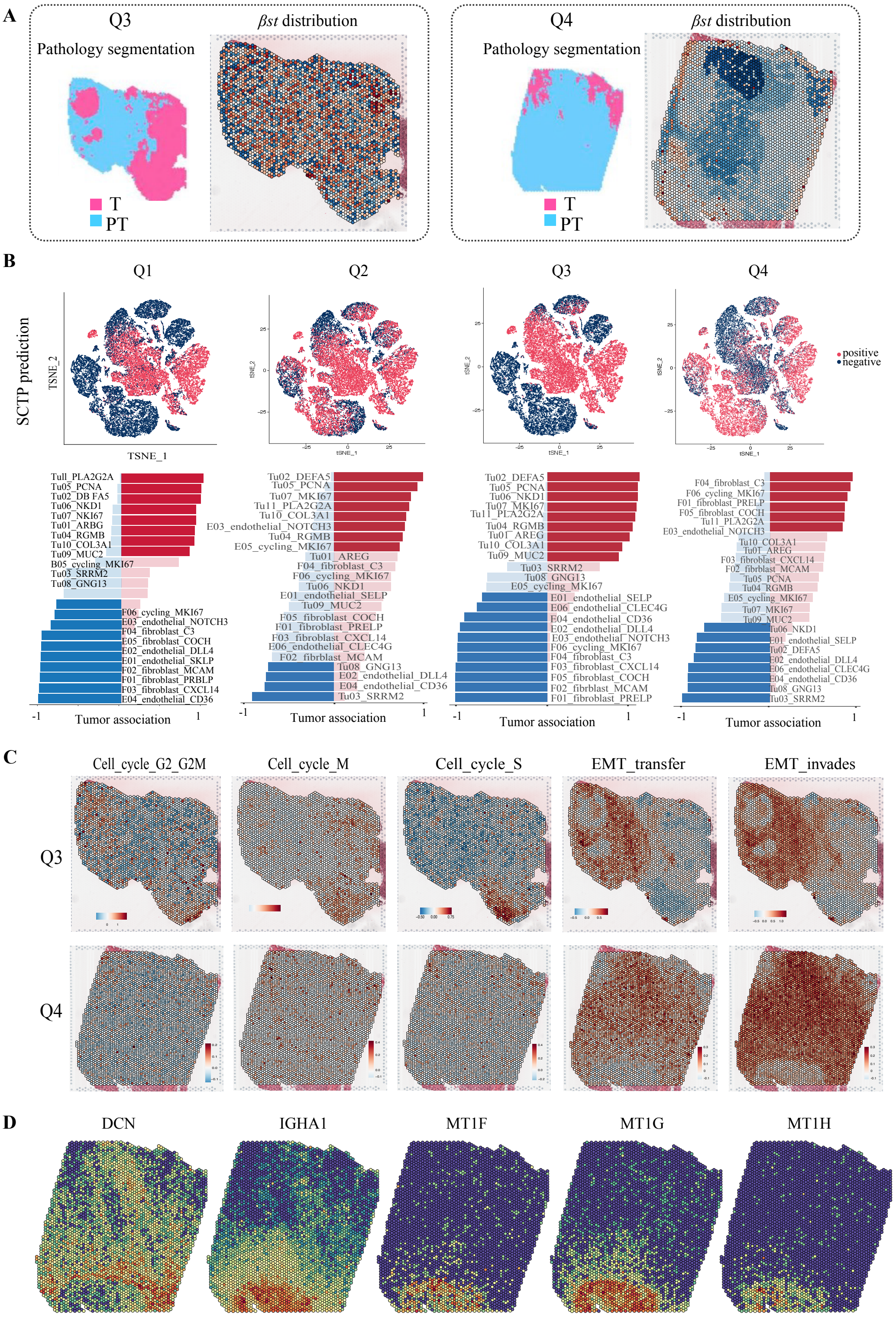


Figure 3: SCTP Model for Colorectal Cancer (SCTP-CRC) and its predictive analysis of tumor microenvironment partitioning. **A**: Segmentation Based on Pathology and Estimation of Latent Space Parameters $\beta_{st}$ for ST data Q3 and Q4, Respectively. **B**: t-SNE visualization of 41,892 non-immune cells, categorized according to the SCTP-CRC model's Predictions on cell malignancy with different ST data (upper panel) and partition of predicted malignant versus non-malignant cells across cell subtypes (lower panel). **C**: Gene expression levels within pathways associated with cell cycle (G2M, M and S phases respectively), Epithelial-Mesenchymal Transition (EMT) transfer, and EMT invasion. D: Feature plots illustrating expression levels of specific genes: Decorin (DCN), Immunoglobulin Heavy Constant Alpha 1 (IGHA1), Metallothionein 1F (MT1F), Metallothionein 1G (MT1G), and Metallothionein 1H (MI1H).

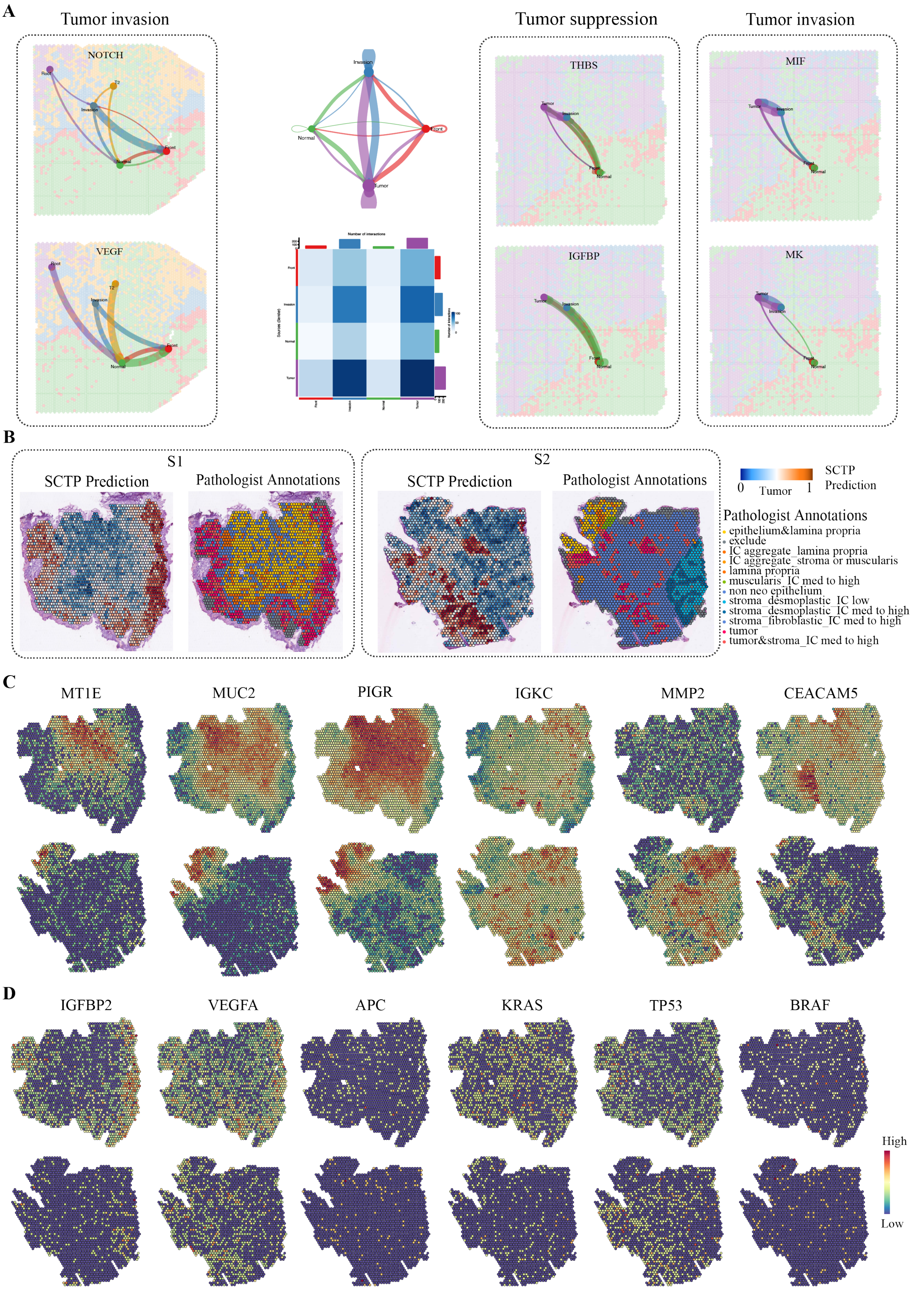


Figure 4: SCTP-CRC accurately predicted tumor area and peri areas in external CRC ST data. **A** Inferred cell-cell communication network visualized by a circle plot and heatmap, displaying the number of interactions between any two cell groups. **B** Predicted tumor and peri-tumor areas with corresponding pathologist annotations for two spatial datasets S1 and S2, respectively. **C** Expression levels of genes MT1E, MUC2, PIGR, IGKC, MMP2, and CEACAM5 in ST samples S1 and S2. **D** Expression levels of genes IGFBP2, VEGFA, APC, KRAS, TP53, BRAF in ST samples S1 and S2.

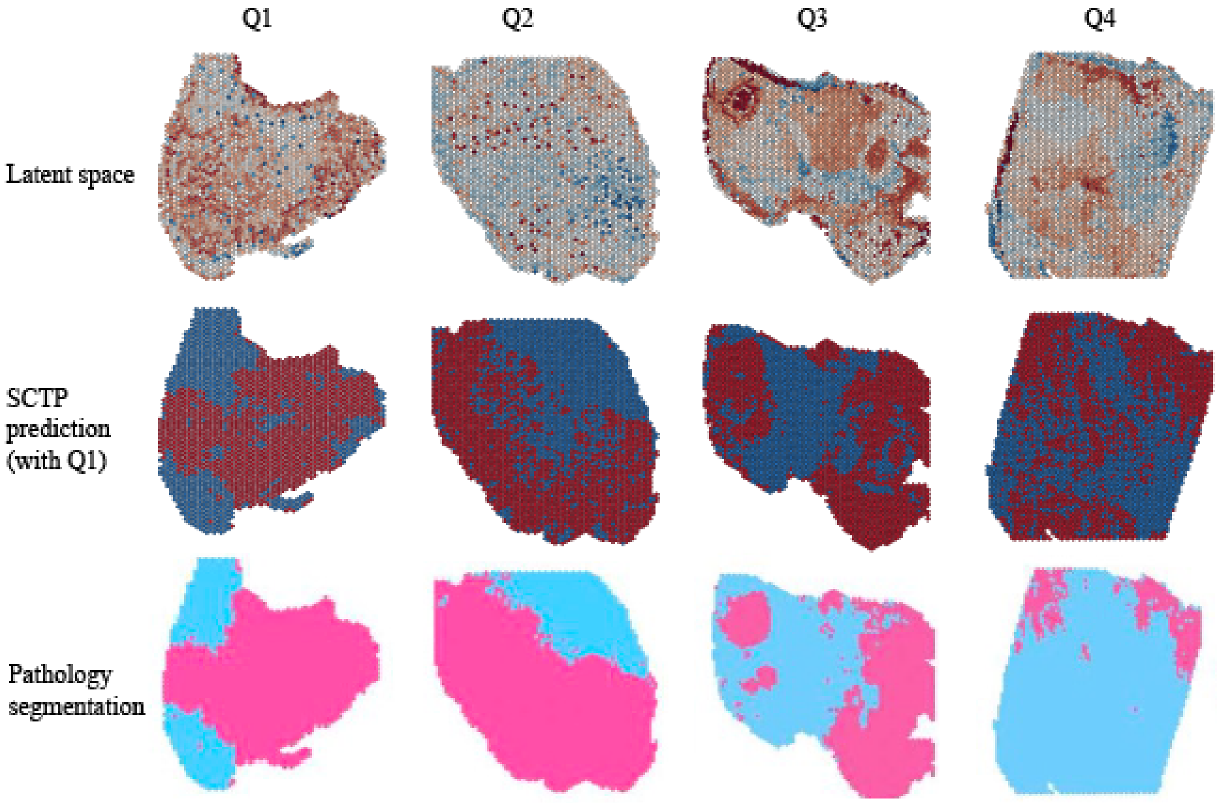


Figure 5: SCTP utilizing single modality in the form of spatial transcriptomic data. Latent space: estimated coefficient $\beta_{st}$ mapped to each spot on the histology image. SCTP prediction: predicted tumor likelihood of each spot based on the estimated coefficient from ST data Q1. Red indicates tumor spots, while blue signifies peri-tumor spots. Pathology segmentation: annotations derived from pathology in the initial study..
